# LLM-PathwayCurator transforms enrichment terms into audit-gated decision-grade claims

**DOI:** 10.64898/2026.02.18.706381

**Authors:** Ken Furudate, Koichi Takahashi

## Abstract

Pathway enrichment of omics data requires careful interpretation, and LLM-PathwayCurator transforms enrichment outputs into auditable, evidence-linked claims with redundancy-highlighting module mapping. Across seven cohorts from The Cancer Genome Atlas (TCGA), it achieved qualified coverage of 0.66 to 0.80 but fell to a range of 0.20 to 0.42 under context swap and 0.20 to 0.30 after supporting-gene dropout due to contract violations or weak gene support. LLM-PathwayCurator generates decision-grade claims with audit-gated abstention to establish a reproducible quality-assurance layer for omics interpretation.

---

Pathway enrichment analysis is a standard approach for interpreting omics data^1^, yet in practice it returns enrichment terms and summary statistics, leaving the endorsement of any particular interpretation to the analyst. Analysts select representative terms from clusters of near-duplicates and subjectively judge interpretation strength, limiting reproducibility. While large language models (LLMs) can assist narrative interpretation^2^, free-text narratives are challenging to reproduce and cannot be rule-audited because they lack verifiable claim-to-evidence links (term identifiers and supporting genes). Consequently, candidate interpretations cannot be systematically audited for evidence-link drift, internal contradiction, context specificity, or fragility under supporting-gene perturbations.

To address this gap, we developed LLM-PathwayCurator, which transforms enrichment interpretations into evidence-linked, audit-gated decisions. LLM-PathwayCurator normalizes enrichment outputs from rank-based methods (e.g., fgsea^3^, which implements the GSEA method^4^) and over-representation analysis (ORA; e.g., Metascape^5^) into a term–gene EvidenceTable that records each enriched term and its supporting genes (**Supplementary Table 1 and 2)**. It applies deterministic supporting-gene perturbations to compute stability scores and factorizes the bipartite graph into modules preserving shared support. The workflow is deterministic by default; in optional LLM-assisted mode, the LLM is confined to proposal-only steps, inspired by a blueprint-first, deterministic workflow design^6^, and enforced by predefined rule-based audit gates. The LLM (i) selects context-consistent representatives using a structured Sample Card (condition, tissue, perturbation, comparison) and (ii) emits schema-bounded JSON claims with resolvable EvidenceTable links (term/module identifiers and a supporting-gene set hash; **Extended Data Fig. 1**). Proposed claims are evaluated by predefined audit gates that assign PASS/ABSTAIN/FAIL. It prioritizes audit-qualified claims using a deterministic utility score integrating evidence strength, stability, and context fit, and summarizes redundancy via a module map. Outputs include a reason-coded audit log supporting risk–coverage aware abstention (**Fig. 1; Supplementary Table 3**).

**Fig. 1.**
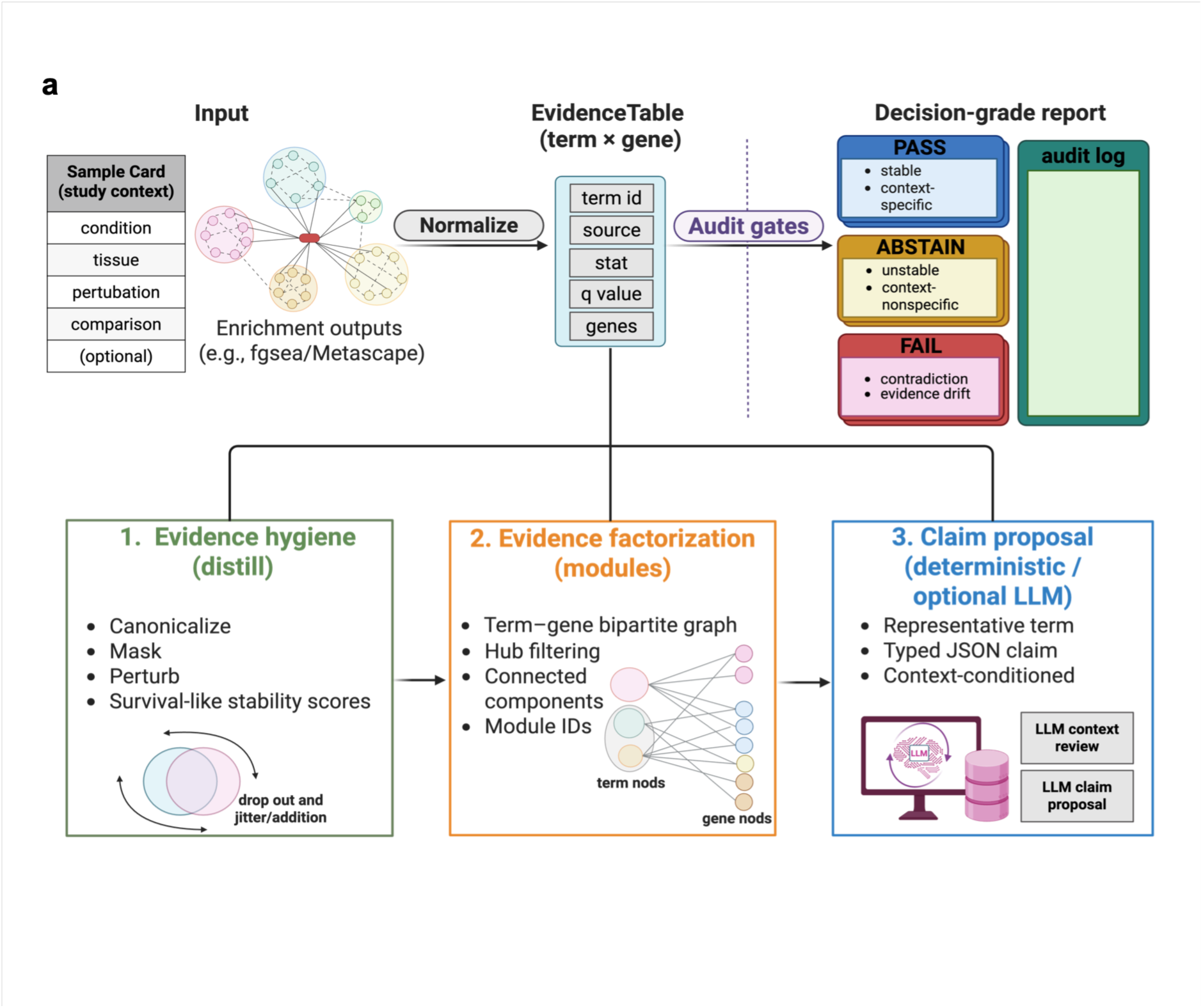
LLM-PathwayCurator transforms enrichment outputs into audited, decision-grade claims. Workflow overview. Enrichment outputs are normalized into a term–gene EvidenceTable conditioned on a Sample Card. Deterministic gene perturbations yield stability scores and the bipartite graph is factorized into evidence modules. The LLM proposes schema-bounded JSON claims with resolvable links; predefined audit gates assign PASS/ABSTAIN/FAIL and emit a reason-coded report

To evaluate decision-grade performance, we applied LLM-PathwayCurator across seven cohorts from The Cancer Genome Atlas^7^ (TCGA; BRCA, HNSC, LUAD, LUSC, OV, SKCM, and UCEC; **Supplementary Table 1**). For each cohort, we first ran fgsea using MSigDB Hallmark gene sets^8^ on a ranked gene list comparing TP53-mutant versus wild-type tumors, and normalized the outputs into a term–gene EvidenceTable (**Supplementary Table 2**). Unless noted otherwise, we used Hallmark gene sets; additional runs with Gene Ontology, Kyoto Encyclopedia of Genes and Genomes and Reactome confirmed that module structure and

ABSTAIN reasons are gene-set-collection dependent (**Extended Data Fig. 2**). Next, we generated *k* = 50 candidate claims per cohort and summarized audit outcomes (PASS/ABSTAIN/FAIL) under three evaluation settings: (i) Proposed (matched Sample Card context), (ii) context swap (shuffled Sample Card context; e.g., BRCA→LUAD), and (iii) evidence-dropout stress (*p* = 0.05). We used a hard-gated audit, in which any triggered gate sets the final decision by precedence (**Supplementary Table 3**). Under Proposed, PASS was 0.66–0.80 (33–40/50) and ABSTAIN was 0.20–0.34 (10–17/50), whereas context swap reduced PASS to 0.20–0.42 (10–21/50) and evidence-dropout reduced PASS to 0.20–0.30 (10–15/50) at *τ* = 0.2, where FAIL was not observed. This reduction is expected: context swap violates the context contract and gene dropout weakens supporting evidence, triggering abstention. Therefore, the audit layer shifts candidates from PASS toward ABSTAIN under context or evidence perturbations rather than endorsing incompatible or fragile interpretations (**Fig. 2a**). Increasing *τ* to 0.9 shifted outcomes toward higher ABSTAIN and lower PASS (e.g., BRCA 0.74→0.70, LUAD 0.74→0.66, OV 0.72→0.62), consistent with *τ* acting as an operating point that trades coverage for conservativeness (**Extended Data Fig. 2a-d; Supplementary Table 4**).

**Fig. 2.**
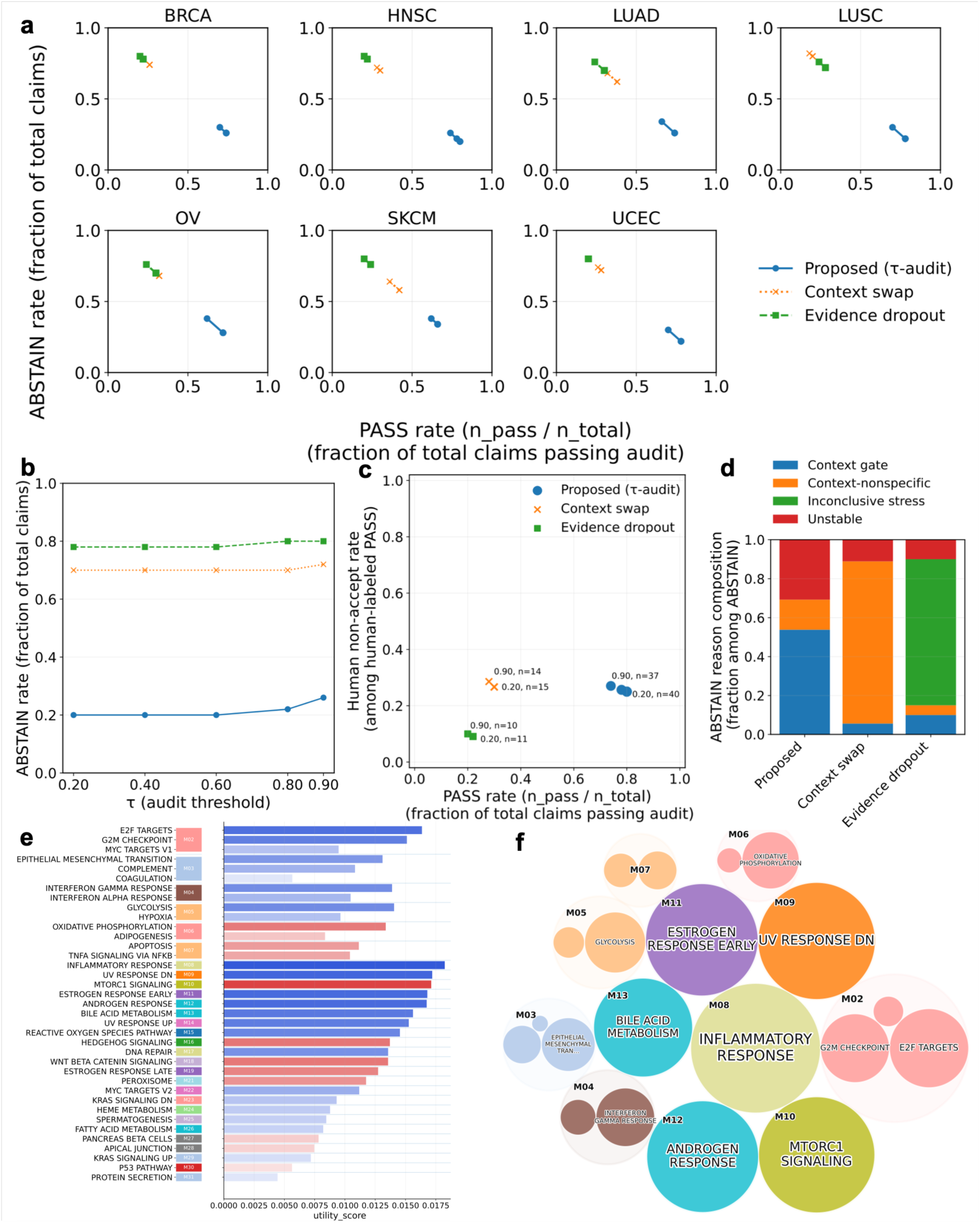
Context validity and robustness of audited claims. **a**, Across TCGA cohorts, *k* = 50 claims were generated and evaluated by predefined audit gates (hard-gated audit) assigning PASS, ABSTAIN, or FAIL. The x-axis shows PASS rate = n_pass/n_total and the y-axis shows ABSTAIN rate = n_abstain/n_total (n_total = 50 per point); FAIL = 0 in this run. Three evaluation settings were compared: Proposed (matched Sample Card context), context swap (shuffled Sample Card context), and evidence-dropout stress (random removal of supporting genes; *p* = 0.05, min_keep = 1). Points correspond to stability thresholds *τ* ∈ {0.2, 0.4, 0.6, 0.8, 0.9}, which tune conservativeness by abstaining on claims with insufficient stability. TCGA cohort sample sizes are listed in **Supplementary Table 1. b–d,** In HNSC, audit-gated abstention reduces human non-accept risk under context and evidence stress. Risk is computed among human-labeled audit-PASS claims (**Methods**; **Supplementary Table 5**). **b**, ABSTAIN rate (n_abstain/50) across *τ* for Proposed, context swap, and evidence-dropout stress (*p* = 0.05, min_keep = 1). **c**, PASS rate (n_pass/50) versus human non-accept risk; point size encodes labeled claims. **d**, ABSTAIN reason composition at *τ* = 0.9, showing stress-specific, rule-attributable reason codes. **e, f,** Utility score–based ordering of audit-PASS claims and redundancy-highlighting module mapping in HNSC (Proposed, *τ* = 0.9). **e**, Audit-PASS claims ordered by a deterministic utility score; bar length indicates utility score and color indicates direction. **f**, Corresponding module map for the same set of claims. Circles represent modules (area proportional to summed utility scores) labeled by the top-ranked term.

To quantify the risk–coverage trade-off, we defined risk as the fraction of human-labeled SHOULD_ABSTAIN or REJECT among audit-PASS claims (**Methods; Supplementary Table 5**) and evaluated the three settings in the HNSC cohort (*k* = 50 claims; hard-gated audit): (i) Proposed, (ii) context swap, and (iii) evidence-dropout stress. Increasing the stability threshold (*τ*) from 0.2 to 0.9 shifted the operating point toward higher abstention under Proposed (PASS 0.80→0.74; ABSTAIN 0.20→0.26), with risk remaining similar (0.25→0.27; 10/40 vs. 10/37 non-acceptable among audit-PASS) as coverage decreased. Under context swap and evidence-dropout stress, coverage decreased further (context swap: PASS 0.30→0.28; evidence-dropout: PASS 0.22→0.20), accompanied by corresponding changes in risk (context swap: 0.27→0.29; evidence-dropout: 0.09→0.10). ABSTAIN reasons shifted accordingly—dominated by context-nonspecific under swap and inconclusive-stress under dropout—consistent with stress-specific, rule-attributable abstention (**Fig. 2b–d**). Using the deterministic utility score defined above, LLM-PathwayCurator produces a reproducible shortlist of audit-PASS claims for triage and reporting (**Fig. 2e**). A companion module map groups claims by shared supporting genes to highlight redundancy and guide representative selection without re-endorsing near-duplicate terms (**Fig. 2f**), providing review- and report-ready outputs rather than supporting specific biological conclusions. Taken together, the audit layer shifted candidates toward stress-specific abstention under context swap, evidence-dropout, or higher *τ*.

To assess the impact of LLM-assisted proposal generation on decision risk, we compared an LLM-assisted setting against a deterministic baseline at a stringent stability threshold (*τ* = 0.8) in the HNSC cohort. Relative to the non-LLM run at the same *τ*, the LLM-assisted run showed lower PASS coverage (0.52 vs. 0.78) and lower human non-accept risk among audit-PASS claims (0.12 vs. 0.26), while shifting more candidates to FAIL (0.46 vs. 0.00; **Extended Data Fig. 3a, b**). All outcomes were mapped to predefined audit reason codes (**Supplementary Table 3**, **6**), with final acceptance determined by rule-based audit gates rather than free-text narratives.

To test generalizability, we applied the same three evaluation settings to the BeatAML2 cohort^9^ (TP53-mutant versus wild-type AML; *k* = 50 claims; hard-gated audit; **Supplementary Table 1**). BeatAML reproduced the TCGA risk–coverage behavior, with high coverage at *τ* = 0.2 (PASS 0.78) and reduced coverage under context swap and dropout (PASS 0.26 and 0.18), while human non-accept risk remained comparable (**Extended Data Fig. 4**). These results support generalizability across independent data sources.

LLM-PathwayCurator improves reproducibility of omics interpretation by transforming plausible but unauditable enrichment interpretations into verifiable, evidence-linked claims. The workflow targets auditable internal consistency rather than biological truth by decoupling proposal from verification via a deterministic audit layer. Unlike significance-ranked term lists or free-text narratives, it provides qualified coverage by passing only context-consistent, evidence-stable claims and abstaining under context shifts or supporting-gene perturbations.

LLM-PathwayCurator can be applied wherever enrichment analysis is performed over curated gene sets, providing a decision-grade quality-assurance layer for omics interpretation.

## Methods

### Overview and study design

LLM-PathwayCurator reframes enrichment interpretation as a decision problem under explicit, auditable constraints. Given an enrichment output and a Sample Card specifying study context (condition, tissue, perturbation, and comparison), the workflow emits schema-bounded JSON claims that link to tool-owned evidence identifiers and that are assigned a discrete decision in {PASS, ABSTAIN, FAIL}. Enrichment outputs from rank-based (e.g., fgsea, RRID:SCR_020938) or ORA methods (e.g., Metascape, RRID:SCR_016620) are normalized into an EvidenceTable that preserves the term–gene relationship. The pipeline comprises (i) deterministic evidence distillation via supporting-gene perturbations (dropout and jitter/addition) to compute per-term survival-like stability scores without re-running enrichment, (ii) evidence factorization by term–gene bipartite modules to retain shared and redundant support, and (iii) constrained claim proposal, where an optional LLM is limited to context review and JSON claim proposal with explicit EvidenceTable links (term identifiers, module identifiers, and supporting-gene set hashes). For ranking and visualization only, we defined a deterministic utility score 𝑈 = 𝑆_evidence_ × 𝑆_’stability_ × 𝑆_context_, where 𝑆_evidence_ is derived from normalized enrichment strength [−log10(q) when available, otherwise ∣ 𝑁𝐸𝑆 ∣or ∣ 𝑠𝑡𝑎𝑡 ∣], 𝑆_’()*#+#(,_ is the aggregated term survival score, and 𝑆_&-%(!.(_is a normalized context-fit proxy; missing components were treated as neutral (1.0), and 𝑈 was not used to assign PASS/ABSTAIN/FAIL. Final decisions are produced by a mechanical audit that enforces evidence-link integrity, a stability threshold (τ), predefined stress tests (context swap and supporting-gene dropout), and intra-run contradiction checks. Spec-level parsing, TSV round-trips, and evidence identity (term_uid, gene-set hashes, and deterministic seeding) are fixed by a centralized contract implementation (**Supplementary Methods**). Evaluation used a two-tier criterion: the primary endpoint was blinded human decision-grade acceptability summarized by risk–coverage analysis, and secondary endpoints were audit reason-code profiles for diagnosis. The workflow was evaluated under three predefined settings—Proposed (matched Sample Card context), context swap, and supporting-gene dropout stress—at fixed operating points defined by *τ*. Evaluation cohorts and comparisons are listed in **Supplementary Table 1**, and the human-labeling protocol is described in **Supplementary Methods** (**Supplementary Table 5)**. Unless otherwise noted, runs used *k* = 50 claims, a hard-gated audit (**Supplementary Table 3**), and MSigDB Hallmark gene sets, with deterministic (non-LLM) proposal generation.

### Inputs and EvidenceTable standardization

All enrichment outputs are normalized (i.e., converted to a common EvidenceTable schema) into a single tool-facing EvidenceTable that preserves the term–gene relationship required for traceability. Each record corresponds to one enriched term and includes a stable term identifier, a numeric enrichment statistic [e.g., NES or −log10(q)], and the term’s supporting genes (field: evidence_genes; leading-edge/core-enrichment genes for rank-based methods, or overlap genes for ORA). EvidenceTable stores supporting genes as a serialized list per term; for downstream graph operations we materialize explicit term–gene edges by expanding this list. Required fields are non-empty term_id, term_name, and evidence_genes, together with a numeric stat. Optional fields (e.g., qval, direction, source) are accepted when present and normalized to a controlled vocabulary; missing qval or direction are carried forward as missing rather than inferred unless a documented deterministic computation is enabled. To support stable joins and audit logs, we construct a tool-owned term_uid from (source, term_id); if source is absent, it is set to a fixed sentinel so that term_uid remains well-defined. We additionally compute a tool-owned supporting-gene set hash from normalized gene tokens (order-invariant), enabling stable claim-to-evidence linkage. Basic validation (required-field checks and numeric coercion; range checks for p/q-values when present) is applied at ingestion. Any deterministic salvage (e.g., coalescing synonymous columns or optional BH-based q-value computation) is recorded in provenance. The EvidenceTable v1 contract and adapter mapping rules—including header normalization, deterministic alias mapping/coalescing, hard validity criteria for row inclusion, and provenance fields (e.g., qval_source)—are fixed in **Supplementary Table 2**.

### Sample Card context specification

Study intent and run configuration are encoded in a Sample Card v1, a tool-facing JSON contract used for claim selection, mechanical audits, and reproducible context stress tests (**Supplementary Methods**). The contract requires four core study-context keys—condition, tissue, perturbation, and comparison—each deterministically canonicalized to a stable string.

Missing, empty, or null (NA) values are replaced with a single sentinel string NA (no null/None semantics). For backward compatibility, disease-like legacy keys (e.g., disease/cancer/tumor) are accepted and deterministically hoisted into condition only when condition is missing or NA, with no other inference. Run configuration (“knobs”, including the stability threshold *τ*) is stored under a top-level extra dictionary that is flattened and canonicalized deterministically (legacy aliases are mapped to current names; unknown keys are preserved for forward compatibility but ignored by audited logic unless explicitly recognized). To prevent configuration drift, k_claims is authoritative only at top level and any occurrences inside extra are removed on load. For context anchoring and stress tests (including context swap), the Sample Card optionally provides deterministic context tokens under a versioned policy (context_token_policy = “ctx_tokens_v1”). Tokens are derived deterministically from an optional text field or from the fallback concatenation, condition|tissue|perturbation|comparison (with NA included literally), and may optionally apply one-hop synonym expansion from a shipped, versioned lexicon with fail-closed semantics. Token provenance (policy identifier and a stable token signature) is recorded for exact reruns.

### Evidence distillation and gene masking

To quantify the robustness of supporting genes without re-running enrichment, the distillation module computes per-term stability scores from deterministic, seed-controlled supporting-gene perturbations. By default, synthetic variants of each term’s supporting-gene set are generated via dropout and jitter/addition (with additions drawn from a fixed global gene pool). A survival-like stability score is defined as the fraction of perturbations whose perturbed supporting-gene sets pass a fixed similarity gate to the original set, with safeguards for small gene sets. All distillation parameters and seeds are recorded as provenance; final PASS/ABSTAIN/FAIL decisions are assigned downstream by the audit layer. Prior to LLM-facing proposal steps, deterministic gene masking is applied to reduce prompt-irrelevant tokens. Masking targets locus-style identifiers (e.g., LOC*/LINC*/Gm*) and lineage-variable TCR/Ig V(D)J patterns, while retaining broad programs (ribosomal, mitochondrial, HLA, cell cycle) by default. Masking uses a symbol-based view for matching but removes tokens from the original supporting-gene lists; in the default implementation, stability scores and the supporting-gene set hash are computed after masking, and masking events (dropped tokens) are recorded as provenance for exact reruns and auditing. Noise definitions were adapted from our marker-centric masking module for cell-type annotation (LLM-scCurator^10^, RRID:SCR_027945) and are cited for provenance. In LLM-PathwayCurator, masking is intentionally conservative and limited to locus-style identifiers and V(D)J-variable prefixes, while broad biological programs are retained by default to avoid systematically altering the evidence supporting audited claims.

### Evidence module factorization

Distilled evidence is factorized into evidence modules by constructing a term–gene bipartite graph from supporting genes. By default, modules are computed as connected components on a term–term graph derived from shared-gene overlap: two terms are linked when they share at least min_shared_genes supporting genes and exceed a minimum Jaccard similarity (jaccard_min; defaults 3 and 0.10). To reduce spurious connectivity driven by high-degree bridge genes, hub genes whose term-degree is strictly greater than a maximum threshold are optionally removed; when an explicit threshold is not provided, it is inferred deterministically from a high quantile of the gene term-degree distribution (default: 0.995 quantile). Overlap graphs can be extremely sparse or can collapse into a dominant component; therefore, the default auto mode applies bounded, deterministic threshold adjustments (relaxing criteria for sparse graphs and tightening criteria when a giant component dominates), and records all effective thresholds and fallbacks as provenance. For very large term sets, connected components on the bipartite graph are used. Module identifiers are content hashes over the unordered sets of terms and genes, yielding stable module IDs independent of traversal order.

### Claim schema and constrained LLM generation

Interpretations are emitted as schema-bounded JSON claims with tool-owned identity and resolvable EvidenceTable links. Each claim references a valid term_uid (optionally a module_id) and carries a supporting-gene set hash (term_gene_set_hash) computed deterministically from the EvidenceTable-derived gene set under the tool’s fixed normalization and capping rules; the tool, not the LLM, fills and verifies identifiers, gene lists, and hashes. For selection and visualization, we computed a deterministic utility score 𝑈 = 𝑆_evidence_ × 𝑆_stability_ × 𝑆_context_, where 𝑆_evidence_derives from normalized enrichment strength [−log10(q) when available, otherwise ∣ 𝑁𝐸𝑆 ∣or ∣ 𝑠𝑡𝑎𝑡 ∣], 𝑆_stability_ is the aggregated term survival score, and 𝑆_context_is a normalized context-fit proxy; missing components were treated as neutral (1.0). This utility score was used only to order and display candidates (e.g., tables/plots), not for assigning PASS/ABSTAIN/FAIL. Free-text narratives are not treated as evidence, and acceptance decisions are never delegated to the LLM. Claim proposal operates in two modes: a deterministic baseline that instantiates claims from the enrichment outputs (using the reported enrichment statistic and q-values as provided), and a constrained LLM mode. In constrained LLM mode, a backend-agnostic model reviews a candidate pool against the Sample Card context under a copy-exact selection regime, selecting term identifiers verbatim from the provided list. For LLM-assisted runs, a local Ollama backend serving llama3.1:8b was used; full backend and inference settings are described in **Supplementary Methods**. A strict validation gate rejects out-of-pool selections, duplicates, and schema-invalid outputs; all evidence-link fields are filled or verified deterministically to preserve stable, audit-grade identity. A reduced audit-log view for the LLM-assisted setting is provided in **Supplementary Table 6**.

### Audit suite and decision rules

Final decisions are produced by a mechanical audit suite with strict precedence (FAIL > ABSTAIN > PASS; **Supplementary Table 3**). The auditor first enforces evidence linkage and identity by resolving referenced terms to unique term_uids and verifying gene_set_hash consistency against the EvidenceTable-derived supporting gene set for the referenced term_uid(s), under the tool’s fixed normalization rules. It then applies stability gating using the survival threshold τ specified by the analysis configuration, and screens for under-supported or hub-dominated evidence to mitigate non-specific bridge effects. Context validity is enforced through a configurable gate; in hard mode, missing context evaluations or WARN (non-specific) context yield abstention under a decision-grade policy. As an additional guardrail, claims that share the same evidence key but assert opposite directions are forced to FAIL (contradiction).

Counterfactual evaluation is defined as internal perturbation only (e.g., context shuffle/swap) and requires no external knowledge. Under counterfactual stress, claim support is expected to weaken or become ambiguous; when the response is ambiguous, the system abstains by default. Context shuffle is treated as a minimal falsification test for context-conditioned selection rather than a semantic gotcha, and is used to verify that selection behavior depends on the Sample Card as intended.

### Calibration and evaluation metrics

Each proposed claim is assigned a discrete audit status in {PASS, ABSTAIN, FAIL}. Coverage is defined as PASS/TOTAL. For internal audit summaries, decided items are defined as PASS+FAIL and audit risk as FAIL/(PASS+FAIL), excluding ABSTAIN from the denominator.

For the external decision-grade endpoint, human non-accept risk among audit-PASS claims is reported, defined as (human-labeled SHOULD_ABSTAIN + human-labeled REJECT) / audit-PASS. Human labels (ACCEPT/SHOULD_ABSTAIN/REJECT) are defined by two independent raters blinded to method variant, audit outcome, and any confidence scores (**Supplementary Methods**; **Supplementary Table 5**) on a fixed labeled subset (HNSC, *n* = 50; BeatAML, *n* = 50), Hard FAIL outcomes are reserved for auditable violations (e.g., contradiction, evidence drift, or contract violations). When human labels are available, confidence scores are optionally calibrated for interpretability using temperature scaling, 𝑝^0^ = 𝜎(logit(𝑝)/𝑇), with 𝑇 > 0 selected by deterministic grid search to minimize binary negative log-likelihood (acceptable = human-labeled ACCEPT; non-acceptable = human-labeled SHOULD_ABSTAIN or human-labeled REJECT). Isotonic regression is also supported as a monotone alternative. Scores are clipped to (𝜀, 1 − 𝜀) prior to any logit transform to avoid numerical instability and can be evaluated on held-out labeled data. Audit-PASS claims are intended to be safe to state as context-bounded observational signature statements under the provided Sample Card; residual non-accept risk is quantified by blinded human labeling. “As written” means without adding substantial qualifiers or rewording to avoid mechanistic/causal implication, or to resolve ambiguity from the provided context/evidence (**Supplementary Table 5**).

### Reporting and provenance

The system generates two complementary report layers to ensure absolute traceability: an audit-grade, machine-readable JSONL artifact (JSONL, report.jsonl) and a human-facing validation bundle (markdown + TSV formats, report.md, audit_log.tsv, distilled.tsv, and, when available, risk_coverage.tsv) that preserve stable keys, decision statuses, and reason codes. Reports include complete provenance of inputs, configuration, effective thresholds, seeds, and software versions, and store sufficient metadata to rerun the pipeline deterministically. Evidence identity is treated as spec-critical; any display-only annotations (e.g., gene symbols) are explicitly non-audited and never change hashing, linkage, or decisions.

### Reproducible figure generation

All data-driven figure panels were generated by scripted pipelines in paper/scripts/, with inputs, outputs, and execution entry points pinned by FIGURE_MAP.csv and per-figure configuration files. Conceptual schematics in **Fig. 1** and **Extended Data Fig. 1** were created with BioRender.com. Online Methods summarizes figure generation at the pipeline-family level and delegates script enumeration and exact command lines to a Supplementary Reproducibility Note. Figure generation is grouped into three reproducible pipeline families corresponding to (i) the primary evaluation figure, (ii) collection and threshold comparison panels, and (iii) BeatAML supplementary analyses. Each family has a single orchestrating entry script that executes end-to-end data preparation, EvidenceTable construction, claim generation, auditing, aggregation, and rendering under fixed seeds and recorded metadata. Pipelines also persist analysis-grade intermediate artifacts (EvidenceTables, audit logs) and run metadata so that panels can be regenerated directly from recorded outputs.

## Data availability

All datasets analyzed in this study are publicly available from their original repositories and publications. TCGA data were accessed via the UCSC Xena platform^11^ (RRID:SCR_018938, https://xenabrowser.net/datapages/). BeatAML^9^ datasets were obtained from the BeatAML2 portal and the associated public data repository (https://biodev.github.io/BeatAML2/). No new datasets were generated in this study.

## Code availability

LLM-PathwayCurator (RRID:SCR_027964) source code, benchmarking scripts, and configuration files are available from GitHub (release v0.1.0; https://github.com/kenflab/LLM-PathwayCurator). A frozen snapshot of the release, together with the source data underlying the figures, is archived on Zenodo^12^. Documentation is available at https://llm-pathwaycurator.readthedocs.io/.

## Acknowledgements

We thank Jeremy Allen (Edanz) for English-language editing of the manuscript.

## Author contributions

K.F. conceived the study. K.F. curated the data; performed formal analysis, validation and investigation; developed the methodology; generated visualizations; and wrote the original draft. K.T. supervised the study and administered the project. K.F. and K.T. reviewed and edited the manuscript.

## Extended Data Figure legends

**Extended Data Fig. 1.**
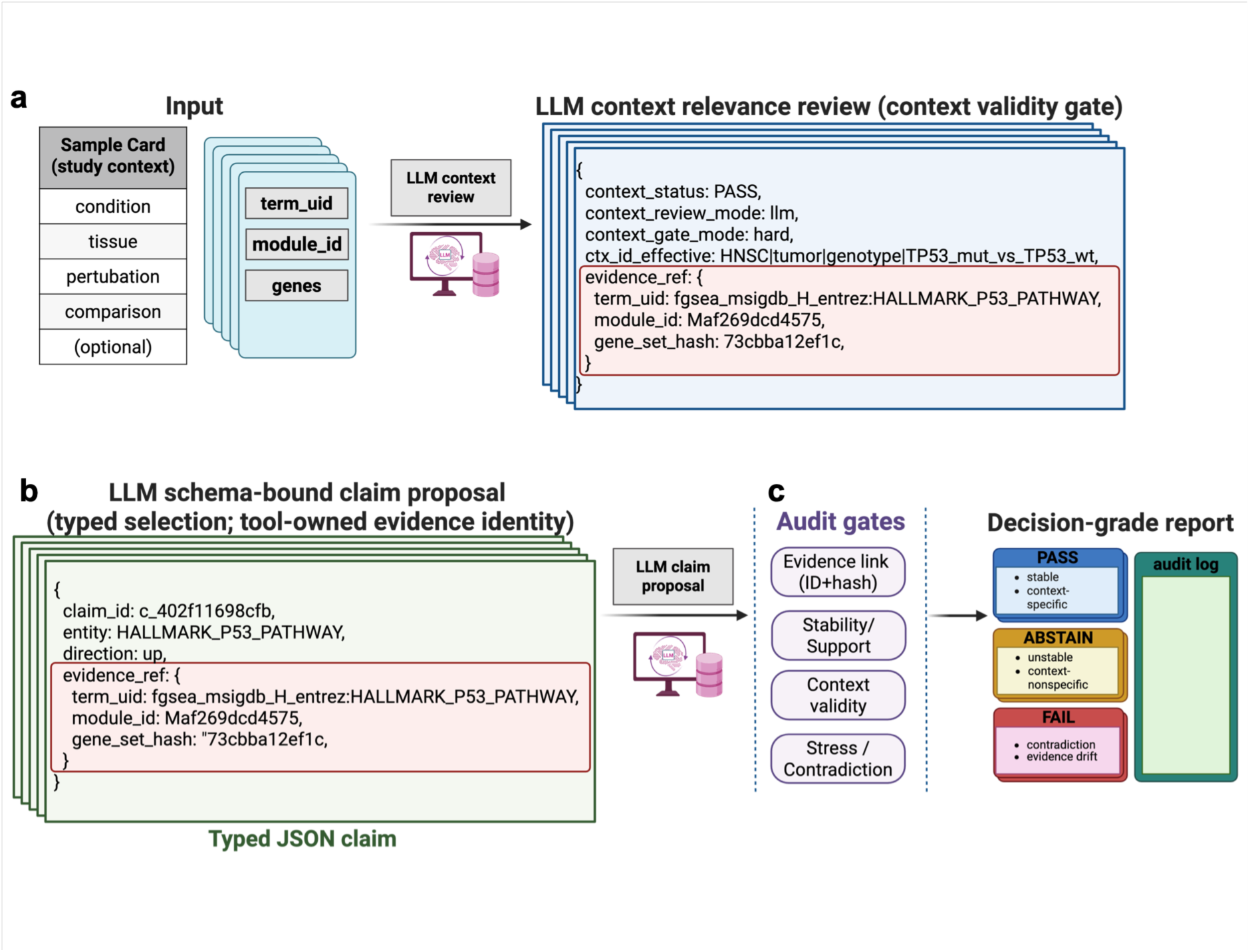
Schema-bounded LLM outputs and predefined rule-based audit gates in LLM-PathwayCurator. This figure specifies the output contracts that make enrichment interpretations machine-auditable: the LLM produces proposals only, and a predefined set of rule-based audit gates assigns the final PASS/ABSTAIN/FAIL decisions with standardized reason codes. **a**, Context relevance review (proposal stage). Given a Sample Card (study context) and evidence-linked identifiers, the LLM returns a structured context review indicating whether the proposed interpretation is context-specific or should be treated as context-nonspecific; this review does not determine acceptance. **b**, Typed claim proposal with evidence-link constraints. The LLM proposes a typed, schema-bounded JSON claim with resolvable links to the EvidenceTable (e.g., term/module identifiers, and a gene-set hash computed from the linked supporting genes). These fields prevent free-text narrative drift and enable deterministic validation of claim–evidence consistency. **c**, Rule-based audit gates and decisions. The audit suite evaluates each proposed claim using (i) evidence-link integrity (IDs/hashes resolve to EvidenceTable entries), (ii) stability/support thresholds (*τ*), (iii) context validity stress tests, and (iv) auditable violations (e.g., internal contradiction or evidence drift when detected). The pipeline outputs decision-grade reports with audit logs that record PASS/ABSTAIN/FAIL and standardized reason codes.

**Extended Data Fig. 2.**
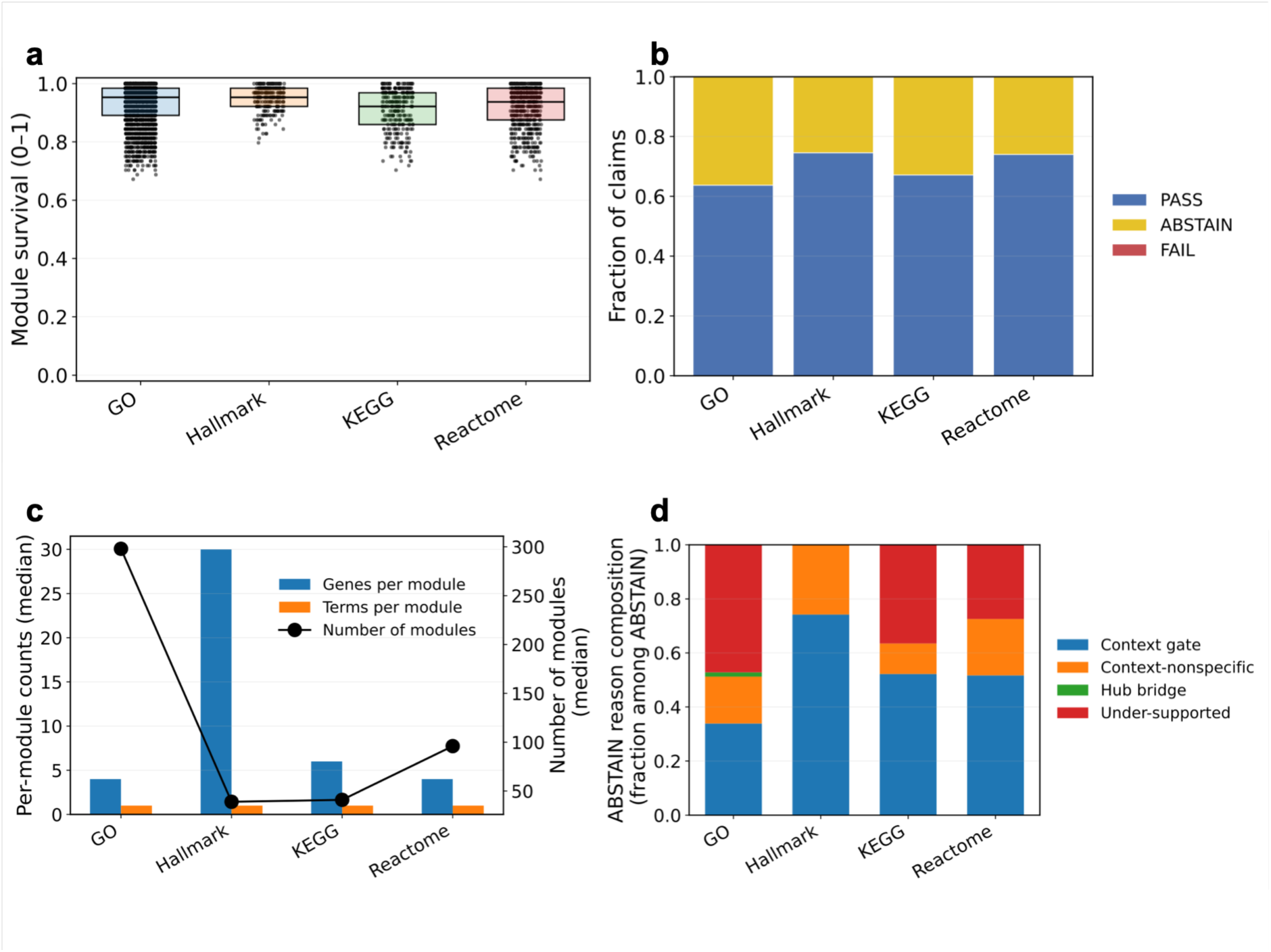
Collection-dependent module structure, stability, and audit outcomes. **a,** Distribution of module stability scores (0–1) across collections under a fixed run setting (*τ* = 0.2; hard-gated audit; Proposed). **b,** Audit outcome composition (PASS/ABSTAIN/FAIL; 100% stacked) aggregated by collection. **c,** Median module structure statistics by collection: genes per module and terms per module (left axis) and the number of modules (right axis). **d,** Composition of ABSTAIN reason codes by collection (top categories; fraction within ABSTAIN), including Context gate, Context-nonspecific, Hub bridge, and Under-supported. All panels use the same benchmark slice and parameterization (PANCAN_TP53_v1; *τ* = 0.2; hard-gated audit; Proposed).

**Extended Data Fig. 3.**
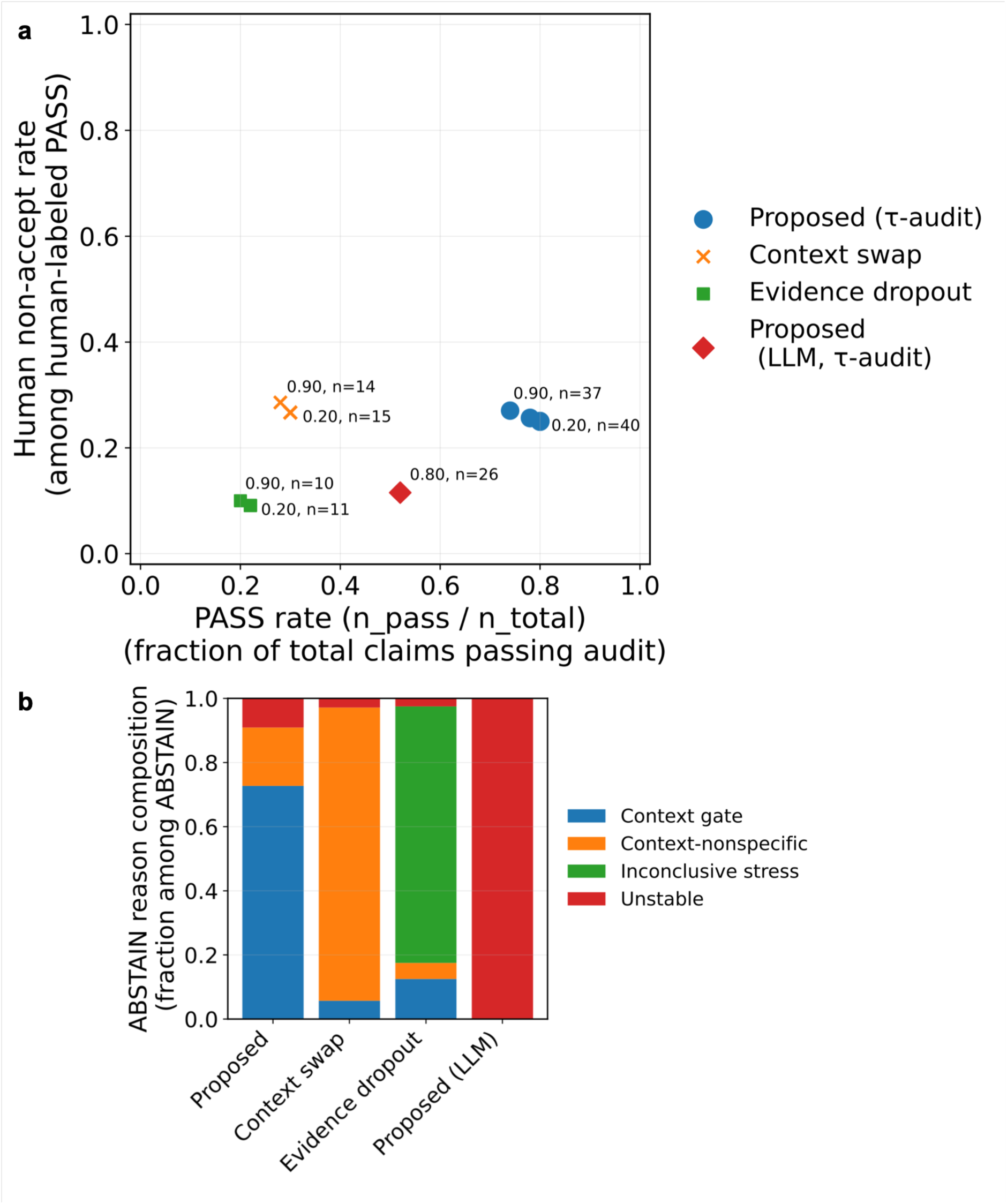
LLM-assisted proposal generation under audit-gated selective abstention in the HNSC cohort. We generated *k* = 50 schema-bounded claims for the HNSC cohort and evaluated all settings using the same hard-gated audit, which assigns each claim to PASS/ABSTAIN/FAIL and records predefined audit reason codes (**Supplementary Table 3**). **a**, PASS coverage (PASS/TOTAL) versus human non-accept risk among human-labeled audit-PASS claims, defined as (human-labeled SHOULD_ABSTAIN + human-labeled REJECT)/audit-PASS (**Methods; Supplementary Table 5**). Points show Proposed (τ-audit), Context swap, and Evidence-dropout stress (*p* = 0.05, min_keep = 1; as in Fig. 2b, c) at *τ* = 0.2 and 0.9, and the LLM-assisted setting under Proposed at *τ* = 0.8. Text annotations report *τ* and n_pass_labeled (point size scales with n_pass_labeled). **b,** ABSTAIN reason-code composition at *τ* = 0.8. Under Proposed, abstentions were primarily assigned to context-gate and unstable codes; under the LLM-assisted setting, abstention was rare and assigned to unstable. Context swap and evidence-dropout stress were dominated by context-nonspecific and inconclusive-stress abstention, respectively.

**Extended Data Fig. 4.**
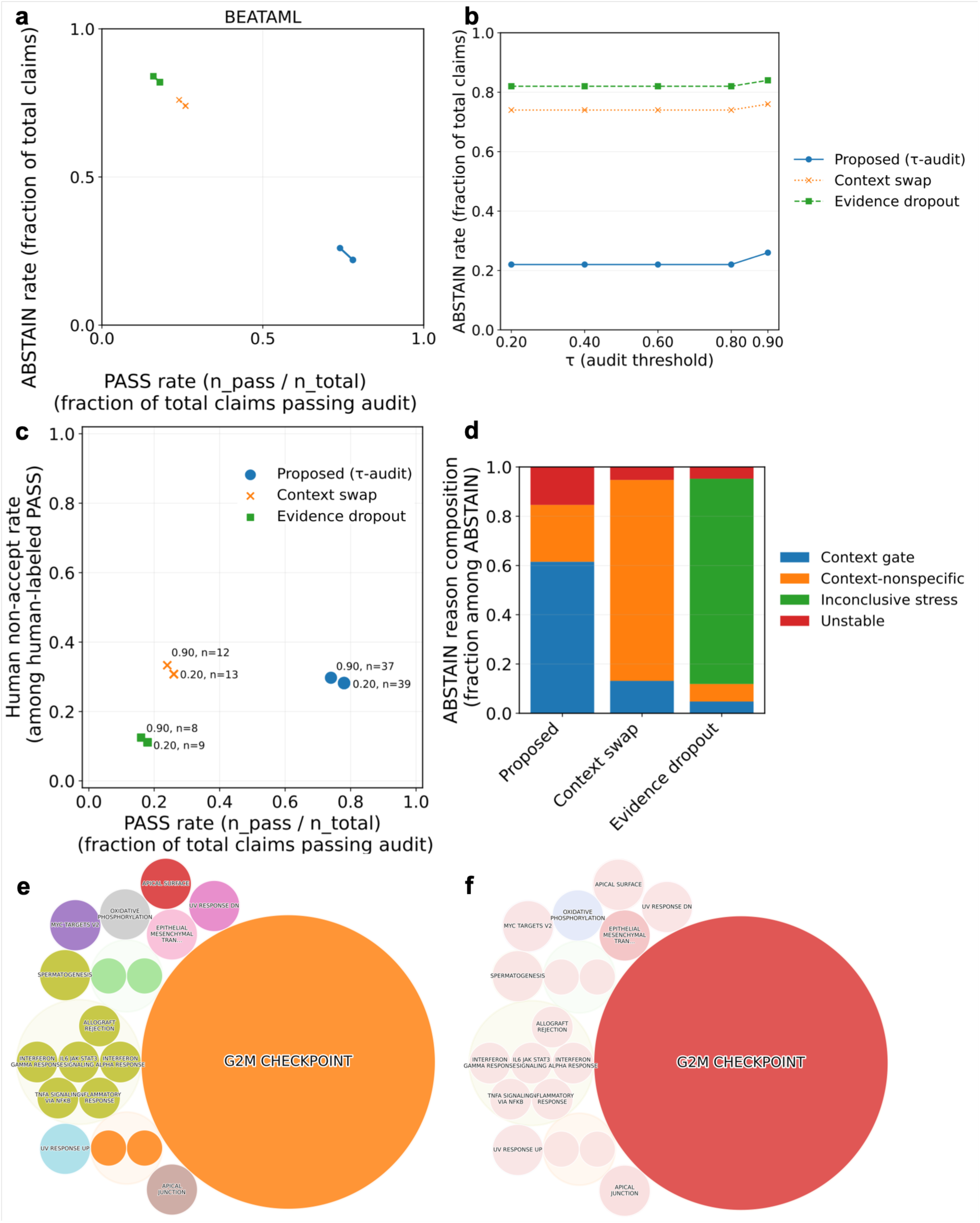
External validation on an independent BeatAML2 cohort. **a-c,** We analyzed the BeatAML2 cohort and generated *k* = 50 typed claims per setting, then applied a hard-gated audit (a predefined set of rule-based audit gates) that assigned each claim to PASS, ABSTAIN, or FAIL (FAIL = 0 in this run). We compared the three settings: Proposed (matched Sample Card context), context swap (Sample Card context keys shuffled to create mismatched context), and evidence-dropout stress (random removal supporting genes; *p* = 0.05, min_keep = 1). Points correspond to stability thresholds *τ* ∈ {0.2, 0.4, 0.6, 0.8, 0.9}, with the x-axis showing PASS rate = n_pass/n_total and the y-axis showing ABSTAIN rate = n_abstain/n_total (n_total = 50 per point). At *τ* = 0.2, Proposed achieved PASS 39/50 (0.78), context swap 13/50 (0.26), and evidence-dropout 9/50 (0.18); at *τ* = 0.9, coverage decreased to 37/50 (0.74), 12/50 (0.24), and 8/50 (0.16), respectively, consistent with conservative tuning via abstention under mismatched Sample Card context and supporting-gene perturbation. BeatAML2 cohort sample sizes are listed in **Supplementary Table 1**. **d,** ABSTAIN reason-code composition at *τ* = 0.9 for Proposed, context swap, and evidence-dropout stress (*p* = 0.05, min_keep = 1). Context swap was dominated by context-nonspecific abstention, whereas evidence-dropout was dominated by inconclusive-stress abstention. **e, f,** Representative redundancy-highlighting module map in BeatAML (Proposed, *τ* = 0.9), ranked by the utility score. Circles represent modules (area proportional to the sum of utility scores) and are labeled with the top-ranked term in each module.

