## Supplementary Information (Methods and Tables) for "LLM-PathwayCurator transforms enrichment terms into audit-gated decision-grade claims"

### Contents

**S1. Supplementary Methods**

**S1.1. Sample Card v1 specification.**

**S1.2. EvidenceTable v1 specification.**

**S1.3. Spec-level parsing and identity policy.**

**S1.4. LLM backend and inference settings (LLM-assisted mode).**

**S1.5. Human labeling protocol (external endpoint: decision-grade acceptability).**

**S2. Supplementary Tables**

**S2.1. Supplementary Table 1 | Evaluation cohorts and comparisons.**

S2.2. Supplementary Table 2 | EvidenceTable specification and adapter mapping

S2.3. Supplementary Table 3 | Audit gates and decision codes (PASS/ABSTAIN/FAIL).

S2.4. Supplementary Table 4 | Risk–coverage summary (primary quantitative table).

S2.5. Supplementary Table 5 | Human labeling protocol and labeled claims.

S2.6. Supplementary Table 6 | Reduced audit-log view (LLM-assisted proposal).

### Supplementary Methods

#### S1.1. Sample Card v1 specification

##### S1.1.1 Overview

Purpose: Encode study intent and run configuration as a tool-facing JSON contract used for claim selection, mechanical audits, and reproducible context stress tests.

Format: UTF-8 JSON object. Token policy is explicitly declared by context_token_policy = "ctx_tokens_v1".

##### S1.1.2. Core context keys (required contract)

Required keys: condition, tissue, perturbation, comparison.

Normalization: Each core value is canonicalized to a stable string (trim + canonical whitespace). Downstream treats canonicalized values as exact tokens.

Missing context: Null, missing, empty, or whitespace-only values are replaced with a single sentinel string NA (no null/None semantics).

Backward compatibility mapping: Disease-like keys (e.g., disease, cancer, tumor) are accepted and deterministically hoisted into condition **only if condition is missing/NA**; no other inference is performed.

#### S1.1.3. Run configuration ("knobs") contract

Location: Run configuration is stored under a top-level extra dictionary (tool-owned, forward-compatible).

Flattening / drift prevention: Nested wrappers (e.g., extra.extra...) are flattened deterministically during load with a fixed collision rule (e.g., inner-most wins). Legacy knob aliases are canonicalized to current key names. Unknown keys are preserved verbatim for forward compatibility, but ignored by audited logic unless explicitly recognized.

#### S1.1.4. k_claims defensive handling (anti-resurrection rule)

Top-level only: k_claims is authoritative only as a top-level key.

Internal mapping: On load, k_claims is mapped to a tool-owned internal field (e.g., k_claims_value) to prevent collisions with extra.

Defensive removal: Any k_claims values (or aliases) found inside extra (including nested wrappers) are removed during load and patch application, preventing resurrection across JSON round-trips or merges.

#### S1.1.5. Deterministic context tokens (ctx_tokens_v1)

Goal: Provide a reproducible token representation for context anchoring and stress tests, not for free-form knowledge injection.

Token source: If context_tokens_text is provided, tokens are derived from that text; otherwise, tokens are derived from the fallback concatenation condition|tissue|perturbation|comparison (with NA included literally).

Tokenization policy (fixed): Unicode normalization (NFKC); lowercasing; deterministic splitting rule; stopword removal (versioned); minimum token length; optional numeric filtering (policy-fixed). Deduplication is order-preserving.

Failure mode: Tokenization is fully deterministic; invalid inputs never trigger external lookups or resource-dependent fallbacks.

**S1.1.6. Optional synonym expansion (fail-closed)**

Optional feature: One-hop synonym expansion can be enabled via a shipped, versioned lexicon.

Fail-closed semantics: If the lexicon is unavailable/malformed or lookup fails, no expansion is applied.

Constraint: Expansion is limited to the shipped lexicon, introduces no external knowledge sources, and uses a deterministic ordering rule for added tokens.

**S1.1.7. Token signature and provenance payload**

Signature: The final token list is summarized by a stable short signature (e.g., sha256(token_string)[:12]), where token_string is a deterministic join of tokens under the policy.

Recorded provenance (recommended): token list (or reproducible reference), signature, policy identifier (ctx_tokens_v1), and effective options (e.g., numeric filtering, synonym expansion on/off, lexicon version/checksum).

Use: Signature and provenance are emitted into audit logs to support exact reruns and traceability.

**S1.1.8. Output invariants (downstream assumptions)**

All four core keys exist and are strings (possibly NA). Knobs are flattened and canonicalized;

unknown keys do not affect audited contracts. k_claims is authoritative only at top-level. ctx_tokens_v1 tokens and signature are deterministic given the Sample Card and the declared policy.

#### S1.2. EvidenceTable v1 specification

EvidenceTable.read_tsv() enforces a tool-facing contract that preserves the term×gene relation. Inputs are read with keep_default_na=False; headers are normalized and mapped through a fixed alias table (ALIASES), with deterministic left-to-right coalescing of duplicate mapped columns recorded in df.attrs. Rows are invalid if required fields are empty, stat is non-numeric, evidence_genes is empty after parsing, or pval/qval falls outside [0,1]; invalid rows are dropped by default (drop_invalid=True) or raised under strict=True. Parsing and canonicalization of gene tokens are spec-owned by _shared (parse_genes, clean_gene_token, join_genes_tsv), and provenance/health summaries are recorded in df.attrs.

#### S1.3. Spec-level parsing and identity policy

To prevent contract drift across adapters, distillation, auditing, TSV round-trips, spec-level parsing, and identity rules are centralized in _shared.py. Token-level missing values are recognized by a fixed NA vocabulary (is_na_token), while scalar missingness is handled separately (is_na_scalar). By convention, “NA” is reserved for missing Sample Card context values, whereas “unknown” is used to fill missing EvidenceTable source fields for stable term identifiers. Evidence gene tokens are parsed conservatively (parse_genes): explicit delimiters (comma/semicolon/pipe) are preferred, bracketed list wrappers are tolerated without evaluation, and whitespace splitting is applied only when all tokens match a gene-like pattern. Tokens are cleaned by a minimal normalizer (clean_gene_token) without forced uppercasing (species/ID-system dependent). For TSV-safe emission, gene lists are joined using a single stable delimiter (GENE_JOIN_DELIM = ";") via join_genes_tsv, preserving input order. Supporting-gene set fingerprints are computed by a set-stable, order-invariant short hash (hash_gene_set_12hex): tokens are conservatively cleaned, de-duplicated, sorted, concatenated under a stable payload format, and hashed to a 12-hex identifier. Stable term identifiers are constructed as term_uid = "<source>:<term_id>" with source defaulting to "unknown" when absent (make_term_uid). Deterministic per-term RNG seeds are derived from (seed, term_uid, term_row_id) using a stable hash function (seed_for_term) to ensure cross-platform reproducibility.

**S1.4 LLM backend and inference settings (LLM-assisted mode).**

LLM-assisted claim proposal and context review were executed using a local Ollama backend serving llama3.1:8b (LLMPATH_BACKEND=ollama; LLMPATH_OLLAMA_MODEL=llama3.1:8b). Context review was performed by the LLM (LLMPATH_CONTEXT_REVIEW_MODE=llm) under hard context gating (LLMPATH_CONTEXT_GATE_MODE=hard). Prompts were restricted to a fixed candidate pool of the top 80 ranked terms (LLMPATH_LLM_TOPN=80; LLMPATH_LLM_MAX_CAND_LINES=80), and selection was enforced under a strict-k regime to require copy-exact term identifiers from the provided list (LLMPATH_LLM_STRICT_K=1). To accommodate local inference latency, request timeouts and escalation were set as follows:

LLMPATH_OLLAMA_READ_TIMEOUT=1500,

LLMPATH_OLLAMA_READ_TIMEOUT_MAX=4000,

LLMPATH_OLLAMA_TIMEOUT_ESCALATIONS=3, and

LLMPATH_OLLAMA_TIMEOUT_FACTOR=2.0.

#### S1.5. Human labeling protocol (external endpoint: decision-grade acceptability)

We evaluated an external endpoint, decision-grade acceptability, by human labeling a fixed subset of proposed claims from two analyses: HNSC ($n$ = 50) and BeatAML ($n$ = 50), for a total of n = 100 claims. Human labels reflect acceptability to state a context-bounded directional enrichment/signature claim under the provided Sample Card, not biological mechanistic truth.

**Label set**. Raters assigned one of three labels (**Supplementary Table 5**): ACCEPT, SHOULD_ABSTAIN, or REJECT. ACCEPT indicates the claim is acceptable to state as a context-bounded directional enrichment/signature observation (not a mechanistic or causal statement). SHOULD_ABSTAIN is reserved for cases where the claim is not safe to communicate as written even under the signature framing (e.g., it would require substantial qualifiers to avoid mechanistic implication, or it is not interpretable from the provided context/evidence without guesswork). REJECT indicates broken/ambiguous evidence linkage or incompatibility with the stated context.

**Materials and standardization**. Each labeled item was presented as a single claim-level record containing a stable claim_id and the minimal fields required for context-bounded, evidence-linked evaluation: condition, entity (term identifier), direction, term_name, the effective evidence module identifier (module_id_effective), and the referenced evidence genes (gene_symbols_str). The corresponding Sample Card context was provided alongside each claim. These rater-facing records are generated deterministically using paper/scripts/fig2_make_labels_template.py from the pipeline's EvidenceTable-derived artifacts and audit outputs; provenance-only fields (e.g., _src) are retained for reproducibility but are not shown during rating.

**Blinding and independence**. Two independent raters labeled all items while blinded to method variant, audit outcome, stability threshold settings, and any confidence scores. Claims were randomized and identified only by stable claim identifiers; raters worked independently without discussion.

**Decision questions and deterministic mapping**. Raters answered three fixed questions (**Supplementary Table 5**): Q1 (evidence linkage pre-screen), Q2 (context-bounded acceptability without overreach), and Q3 (withhold because not safe to communicate as written). Labels were assigned deterministically as: if Q1=No → REJECT; else if Q2=No → REJECT; else if Q3=Yes → SHOULD_ABSTAIN; otherwise → ACCEPT.

**Final label and agreement**. We report per-rater labels and define a conservative final label per claim as the more cautious of the two ratings under the ordering REJECT > SHOULD_ABSTAIN > ACCEPT. Inter-rater agreement was quantified using Cohen's κ for the three-class labels and a binary collapsed endpoint (non-acceptable = SHOULD_ABSTAIN or REJECT).

### S2. Supplementary Tables

#### S2.1. Supplementary Table 1 | Evaluation cohorts and comparisons

This table pins the evaluation cohorts and TP53-mut versus TP53-wt comparisons used to assess decision-grade performance of an interpretation quality-assurance framework; it is not intended to support biological claims. For TCGA cohorts, counts reflect primary-like tumors included in the analysis and stratified by TP53 mutation status. For the BeatAML cohort, eligible RNA sequencing (RNA-seq) samples were restricted to subjects marked for manuscript analyses with RNA-seq available, and TP53-mut was defined by protein-altering TP53 variants in whole-exome sequencing (WES) mapped to RNA-seq samples. All cohorts and counts are fully determined by the pinned group files generated by the reproducible figure pipelines (pipeline IDs shown).

| cohort_id | condition | comparison | n_mut | n_wt | inclusion_rule | pipeline_id |
|---|---|---|---|---|---|---|
| TCGA | BRCA | TP53-mut vs TP53-wt | 264 | 837 | Primary-like tumors only (sample_type_id _ {1,3}) | PANCAN_TP53_v1 |
| TCGA | HNSC | TP53-mut vs TP53-wt | 357 | 171 | Primary-like tumors only (sample_type_id _ {1,3}) | PANCAN_TP53_v1 |
| TCGA | LUAD | TP53-mut vs TP53-wt | 259 | 261 | Primary-like tumors only (sample_type_id _ {1,3}) | PANCAN_TP53_v1 |
| TCGA | LUSC | TP53-mut vs TP53-wt | 401 | 105 | Primary-like tumors only (sample_type_id _ {1,3}) | PANCAN_TP53_v1 |
| TCGA | OV | TP53-mut vs TP53-wt | 57 | 535 | Primary-like tumors only (sample_type_id _ {1,3}) | PANCAN_TP53_v1 |
| TCGA | SKCM | TP53-mut vs TP53-wt | 9 | 95 | Primary-like tumors only (sample_type_id _ {1,3}) | PANCAN_TP53_v1 |
| TCGA | UCEC | TP53-mut vs TP53-wt | 178 | 369 | Primary-like tumors only (sample_type_id _ {1,3}) | PANCAN_TP53_v1 |

| | | | | | | |
|---|---|---|---|---|---|---|
| BeatAML | AML | TP53-mut vs TP53-wt | 58 | 552 | Clinical cohort; RNA-seq; TP53-mut from WES. | BEATAML_TP53_v1 |

TCGA inputs were retrieved from UCSC Xena; BeatAML from the pinned release files. BeatAML inclusion: RNA-seq samples flagged for analysis in the clinical file; TP53-mut defined by protein-altering TP53 variants from WES mapped to RNA-seq samples.

**Abbreviations:** TCGA, The Cancer Genome Atlas; BRCA, breast invasive carcinoma; HNSC, head and neck squamous cell carcinoma; LUAD, lung adenocarcinoma; LUSC, lung squamous cell carcinoma; OV, ovarian serous cystadenocarcinoma; SKCM, skin cutaneous melanoma; UCEC, uterine corpus endometrial carcinoma; BeatAML, Beat acute myeloid leukemia; mutant, mut; wild-type, wt; WES, whole-exome sequencing; RNA-seq, RNA sequencing; n_mut, number of TP53-mut samples; n_wt, number of TP53-wt samples.

### S2.2. Supplementary Table 2 | EvidenceTable specification and adapter mapping

This table fixes the tool-facing EvidenceTable v1 contract and adapter mapping rules that normalize heterogeneous enrichment outputs into a single auditable representation preserving the term×gene relation. Provenance and health summaries are recorded in table metadata (df.attrs) and output fields (e.g., qval_source).

#### Part A | EvidenceTable v1 specification (contract)

**Global rules.** Input tables are read with keep_default_na=False to prevent unintended NA coercion of gene tokens. Column headers are normalized (trim; BOM removal; spaces/dashes→underscore; collapse repeated underscores; lowercase) and mapped through a fixed alias table (ALIASES). When multiple input columns map to the same contract name, they are coalesced deterministically left-to-right by filling only empty cells; the action is recorded in df.attrs["coalesced_duplicate_columns"].

**Row validity (hard contract).** A row is invalid if required identifiers are empty (term_id or term_name), stat is non-numeric, evidence_genes parses to an empty list, or provided pval/qval is outside [0,1]. Invalid rows are dropped by default (drop_invalid=True); strict=True raises on the first invalid row.

**Part A table.** Per-column rules (type, allowed values, NA policy, canonicalization) are fixed below

| column_name | requirement | type (normalized) | allowed values | NA policy | canonicalization_rule (deterministic) |
|---|---|---|---|---|---|
| term_id | Core | string | non-empty | NA-like tokens become empty _ row invalid (unless salvaged) | strip; NA tokens detected by _shared.is_na_token are treated as missing |
| term_name | Core | string | non-empty | NA-like tokens become empty _ row invalid (unless salvaged) | strip; NA tokens treated as missing |
| stat | Core | float | finite numeric | non-numeric _ row invalid | pd.to_numeric(errors="coerce") |
| evidence_genes | Core | list[str] (internal); TSV string on write | ≥1 gene token | empty list _ row invalid | parsed by _shared.parse_genes; written by _shared.join_genes_tsv |
| source | optional (autofill) | string | non-empty | missing/empty _ "unknown" | required-string cleaning; empty replaced with "unknown" |

| | | | | | |
|---|---|---|---|---|---|
| direction | optional (autofill) | string | normalized vocabulary (e.g., up, down, na) | missing _ "na" | normalized by _shared.normalize_direction |
| qval | optional (autofill) | float | in [0,1] when present | may be NA | numeric coercion; if missing but pval present, BH fill may be applied (see below) |
| pval | optional (autofill) | float | in [0,1] when present | may be NA | numeric coercion; range-checked to prevent logP mis-mapping |
| group_id | optional (autofill) | string | any | may be NA | created if missing; no normalization beyond scalar handling |
| is_summary | optional (autofill) | bool | true/false | missing/NA _ false | parsed from common truthy strings (1/true/yes/...) |
| evidence_genes_str | internal | string | N/A | N/A | derived from parsed evidence_genes via _shared.join_genes_tsv; used for TSV-safe emission |
| is_valid / invalid_reason | internal | bool / string | N/A | N/A | produced by the schema gate for auditing and (optional) filtering |

| | | | | | |
|---|---|---|---|---|---|
| term_id_salvaged / term_name_salvaged | internal | bool | N/A | N/A | salvage flags (see "salvage" note below) |
| qval_source | internal | string | qval, bh(pval), missing | N/A | provenance of q-values (input vs BH-filled vs missing) |

### Part B | Adapter mapping rules (input_source → EvidenceTable)

**Adapter rule (hard requirement).**
Adapters MUST preserve the term×gene relation: each output row must correspond to a term and carry a non-empty gene list that represents the evidence used for that term in the upstream tool (e.g., overlap genes for ORA, leading-edge/core enrichment genes for fgsea). Output TSVs are schema-gated by EvidenceTable.read_tsv.

#### B1. fgsea (rank-based) mapping (reference: adapters/fgsea.py)

| required input columns | mapping | notes |
|---|---|---|
| fgsea result table (TSV/CSV/whitespace) with pathway, leadingEdge, and at least one of {NES, ES}; optional {padj, pval} | term_id ← pathway (default term_id_mode="raw") ; term_name ← pathway ; source ← config.source_name (default "fgsea") ; stat ← NES if present else ES ; qval ← padj only ; direction ← sign(NES) if present else sign(stat) ; evidence_genes ← leadingEdge parsed by _shared.parse_genes ; TSV join=';' ; pval stored separately when present | drops rows with missing qval by default (drop_na_qval=True); output sorted by qval asc, \|stat\| desc (no explicit 3rd-key tie-break); raises on empty pathway, empty genes (require_genes=True), or non-numeric stat. |

#### B2. Metascape (ORA) mapping (reference: adapters/metascape.py)

| required inputs | mapping | notes |
|---|---|---|

| | | |
|---|---|---|
| deg_ranking.tsv with gene, score; gene IDs matched to MSigDB sets | term_id ← Term (GO#### → GO:####) ; term_name ← Description ; source ← config.source_name (default "metascape") ; stat ← abs(Log(q-value)) if numeric else abs(LogP) ; qval ← reconstructed from Log(q-value) via sign inference (log10(q) if <=0 else -log10(q)) ; direction ← "na" ; evidence_genes ← Symbols (preferred) else Genes ; parsed by _shared.parse_genes ; TSV join=';' ; group_id ← GroupID ; is_summary ← GroupID.endswith("_Summary") | summary rows excluded by default (include_summary=False); rows with missing reconstructed qval dropped by default (drop_na_qval=True); genes are cleaned/split/dedup first-seen (no sorting stated in code); raises on empty Term/Description, empty genes, or non-numeric Log(q-value)/LogP. |

### S2.3. Supplementary Table 3 | Audit gates and decision codes (PASS/ABSTAIN/FAIL)

Final decisions are produced by a mechanical audit suite with strict precedence (FAIL > ABSTAIN > PASS). Each gate is defined by a machine-checkable trigger, referenced fields (EvidenceTable and claim JSON), a stable reason_code, and enforcement (hard or note).

| gate_id | gate_name | decision | trigger_short | trigger_details | fields_codes | reason_code | enforcement |
|---|---|---|---|---|---|---|---|
| P0 | all gates satisfied | PASS | No FAIL/ABSTAIN | No FAIL gate triggered AND no ABSTAIN gate triggered | O1 | ok | hard |
| F1 | evidence identity drift | FAIL | Evidence link mismatch | term_ids unresolvable/ambiguous OR gene_set_hash mismatch OR (gene_ids present AND overlap < min_overlap) | C1,C2,C3,D1,D2,S1 | evidence_drift | hard |
| F2 | schema violation | FAIL | Schema/evidence-link invalid | claim JSON violates schema OR required evidence links are missing/unresolvable | C6,X1 | schema_violation | hard |
| F3 | contradiction | FAIL | Incompatible directions | two claims share the same evidence key but assert incompatible directions (forced failure) | C4,C5 | contradiction | hard |
| F4 | context hard-fail | FAIL | Context gate FAIL | context gate returns FAIL under configured policy | A1,S2 | context_fail | hard |
| A1 | unstable support | ABSTAIN | Stability < τ | stability score below threshold τ | D3,S3 | unstable | hard |
| A2 | missing survival | ABSTAIN | Missing stability | survival/stability fields required for gating are absent | D3 | missing_survival | hard |
| A3 | missing evidence genes | ABSTAIN | Missing support genes | gene_set_hash missing/invalid OR EvidenceTable support genes unavailable for referenced term_uid(s) | C2,D2,S4 | missing_evidence_genes | hard |
| A4 | context missing | ABSTAIN | Context not evaluated | Context missing: not evaluated or status missing/unknown; swap strictness also abstains | C7,C8,S2,S5,R1 | context_missing | hard |
| A5 | context non-specific | ABSTAIN | Context non-specific | Context evaluated but WARN (non-specific); swap strictness treats WARN as abstention | C7,C8,C10,S2,S5,R1 | context_nonspecific | hard |
| A6 | under-supported | ABSTAIN | Union genes < min | Under-supported: \|union evidence genes\| < min_union_genes | D2,S6 | under_supported | hard |

| | | | | | | | |
|---|---|---|---|---|---|---|---|
| A7 | hub bridge | ABSTAIN | Hub bridge high | hub fraction in union evidence genes >= hub_frac_thr (distilled.evidence_genes -> gene degree) | D2,S7,S8 | hub_bridge | hard |
| A8 | inconclusive stress | ABSTAIN | Stress inconclusive | (stress_gate_mode==hard AND stress missing/non-PASS) OR internal stress probe triggers fail-closed stress columns (stress_* / contradiction_*) | S9 | inconclusive_stress | hard |

**fields_codes**:
O1 = audit outputs: decision, reason_code
C1 = claim.term_ids
C2 = claim.gene_set_hash
C3 = claim.gene_ids
C4 = claim.evidence_key
C5 = claim.direction
C6 = claim.* required fields (schema-bounded)
C7 = claim.context_evaluated
C8 = claim.context_status
C10 = context fallbacks: context_review_* / legacy context_* / context_score
D1 = distilled.term_uid
D2 = distilled.evidence_genes(_str) / distilled.evidence_genes_str
D3 = distilled.term_survival_agg / distilled.*survival* fields
A1 = audit.context_status (or equivalent)
S1 = SampleCard.extra.audit_min_gene_overlap (or card.audit_min_gene_overlap)
S2 = SampleCard.extra.context_gate_mode
S3 = SampleCard.extra.audit_tau
S4 = SampleCard.extra.strict_evidence_check
S5 = SampleCard.extra.context_swap_strict_mode
S6 = SampleCard.extra.min_union_genes (or card.min_union_genes)
S7 = SampleCard.extra.hub_term_degree
S8 = SampleCard.extra.hub_frac_thr
S9 = SampleCard.extra.stress_gate_mode (+ probe knobs)
R1 = row.context_swap_active
X1 = schema validation status

**S2.4. Supplementary Table 4 | Coverage and abstention summary across TCGA cohorts (related to Fig. 2)**

We report PASS coverage and ABSTAIN rate across cohorts and $\tau$. No FAIL outcomes were observed, so selective_risk was 0 throughout.

| condition | method | tau ($\tau$) | coverage_pass | abstain_rate_total | n_total | n_pass | n_abstain |
|---|---|---|---|---|---|---|---|
| BRCA | Proposed | 0.2 | 0.74 | 0.26 | 50 | 37 | 13 |
| BRCA | Proposed | 0.4 | 0.74 | 0.26 | 50 | 37 | 13 |
| BRCA | Proposed | 0.6 | 0.74 | 0.26 | 50 | 37 | 13 |
| BRCA | Proposed | 0.8 | 0.74 | 0.26 | 50 | 37 | 13 |
| BRCA | Proposed | 0.9 | 0.7 | 0.3 | 50 | 35 | 15 |
| BRCA | context_swap | 0.2 | 0.26 | 0.74 | 50 | 13 | 37 |
| BRCA | context_swap | 0.4 | 0.26 | 0.74 | 50 | 13 | 37 |
| BRCA | context_swap | 0.6 | 0.26 | 0.74 | 50 | 13 | 37 |
| BRCA | context_swap | 0.8 | 0.26 | 0.74 | 50 | 13 | 37 |
| BRCA | context_swap | 0.9 | 0.26 | 0.74 | 50 | 13 | 37 |
| BRCA | Evidence dropout | 0.2 | 0.22 | 0.78 | 50 | 11 | 39 |
| BRCA | Evidence dropout | 0.4 | 0.22 | 0.78 | 50 | 11 | 39 |
| BRCA | Evidence dropout | 0.6 | 0.22 | 0.78 | 50 | 11 | 39 |
| BRCA | Evidence dropout | 0.8 | 0.22 | 0.78 | 50 | 11 | 39 |
| BRCA | Evidence dropout | 0.9 | 0.2 | 0.8 | 50 | 10 | 40 |
| HNSC | Proposed | 0.2 | 0.8 | 0.2 | 50 | 40 | 10 |
| HNSC | Proposed | 0.4 | 0.8 | 0.2 | 50 | 40 | 10 |
| HNSC | Proposed | 0.6 | 0.8 | 0.2 | 50 | 40 | 10 |
| HNSC | Proposed | 0.8 | 0.78 | 0.22 | 50 | 39 | 11 |
| HNSC | Proposed | 0.9 | 0.74 | 0.26 | 50 | 37 | 13 |
| HNSC | context_swap | 0.2 | 0.3 | 0.7 | 50 | 15 | 35 |
| HNSC | context_swap | 0.4 | 0.3 | 0.7 | 50 | 15 | 35 |
| HNSC | context_swap | 0.6 | 0.3 | 0.7 | 50 | 15 | 35 |
| HNSC | context_swap | 0.8 | 0.3 | 0.7 | 50 | 15 | 35 |
| HNSC | context_swap | 0.9 | 0.28 | 0.72 | 50 | 14 | 36 |

| | | | | | | | |
|---|---|---|---|---|---|---|---|
| HNSC | Evidence dropout | 0.2 | 0.22 | 0.78 | 50 | 11 | 39 |
| HNSC | Evidence dropout | 0.4 | 0.22 | 0.78 | 50 | 11 | 39 |
| HNSC | Evidence dropout | 0.6 | 0.22 | 0.78 | 50 | 11 | 39 |
| HNSC | Evidence dropout | 0.8 | 0.2 | 0.8 | 50 | 10 | 40 |
| HNSC | Evidence dropout | 0.9 | 0.2 | 0.8 | 50 | 10 | 40 |
| LUAD | Proposed | 0.2 | 0.74 | 0.26 | 50 | 37 | 13 |
| LUAD | Proposed | 0.4 | 0.74 | 0.26 | 50 | 37 | 13 |
| LUAD | Proposed | 0.6 | 0.74 | 0.26 | 50 | 37 | 13 |
| LUAD | Proposed | 0.8 | 0.74 | 0.26 | 50 | 37 | 13 |
| LUAD | Proposed | 0.9 | 0.66 | 0.34 | 50 | 33 | 17 |
| LUAD | context_swap | 0.2 | 0.38 | 0.62 | 50 | 19 | 31 |
| LUAD | context_swap | 0.4 | 0.38 | 0.62 | 50 | 19 | 31 |
| LUAD | context_swap | 0.6 | 0.38 | 0.62 | 50 | 19 | 31 |
| LUAD | context_swap | 0.8 | 0.38 | 0.62 | 50 | 19 | 31 |
| LUAD | context_swap | 0.9 | 0.32 | 0.68 | 50 | 16 | 34 |
| LUAD | Evidence dropout | 0.2 | 0.3 | 0.7 | 50 | 15 | 35 |
| LUAD | Evidence dropout | 0.4 | 0.3 | 0.7 | 50 | 15 | 35 |
| LUAD | Evidence dropout | 0.6 | 0.3 | 0.7 | 50 | 15 | 35 |
| LUAD | Evidence dropout | 0.8 | 0.3 | 0.7 | 50 | 15 | 35 |
| LUAD | Evidence dropout | 0.9 | 0.24 | 0.76 | 50 | 12 | 38 |
| LUSC | Proposed | 0.2 | 0.78 | 0.22 | 50 | 39 | 11 |
| LUSC | Proposed | 0.4 | 0.78 | 0.22 | 50 | 39 | 11 |
| LUSC | Proposed | 0.6 | 0.78 | 0.22 | 50 | 39 | 11 |
| LUSC | Proposed | 0.8 | 0.78 | 0.22 | 50 | 39 | 11 |
| LUSC | Proposed | 0.9 | 0.7 | 0.3 | 50 | 35 | 15 |
| LUSC | context_swap | 0.2 | 0.2 | 0.8 | 50 | 10 | 40 |
| LUSC | context_swap | 0.4 | 0.2 | 0.8 | 50 | 10 | 40 |
| LUSC | context_swap | 0.6 | 0.2 | 0.8 | 50 | 10 | 40 |

| | | | | | | | |
|---|---|---|---|---|---|---|---|
| LUSC | context_swap | 0.8 | 0.2 | 0.8 | 50 | 10 | 40 |
| LUSC | context_swap | 0.9 | 0.18 | 0.82 | 50 | 9 | 41 |
| LUSC | Evidence dropout | 0.2 | 0.28 | 0.72 | 50 | 14 | 36 |
| LUSC | Evidence dropout | 0.4 | 0.28 | 0.72 | 50 | 14 | 36 |
| LUSC | Evidence dropout | 0.6 | 0.28 | 0.72 | 50 | 14 | 36 |
| LUSC | Evidence dropout | 0.8 | 0.28 | 0.72 | 50 | 14 | 36 |
| LUSC | Evidence dropout | 0.9 | 0.24 | 0.76 | 50 | 12 | 38 |
| OV | Proposed | 0.2 | 0.72 | 0.28 | 50 | 36 | 14 |
| OV | Proposed | 0.4 | 0.72 | 0.28 | 50 | 36 | 14 |
| OV | Proposed | 0.6 | 0.72 | 0.28 | 50 | 36 | 14 |
| OV | Proposed | 0.8 | 0.72 | 0.28 | 50 | 36 | 14 |
| OV | Proposed | 0.9 | 0.62 | 0.38 | 50 | 31 | 19 |
| OV | context_swap | 0.2 | 0.32 | 0.68 | 50 | 16 | 34 |
| OV | context_swap | 0.4 | 0.32 | 0.68 | 50 | 16 | 34 |
| OV | context_swap | 0.6 | 0.32 | 0.68 | 50 | 16 | 34 |
| OV | context_swap | 0.8 | 0.32 | 0.68 | 50 | 16 | 34 |
| OV | context_swap | 0.9 | 0.32 | 0.68 | 50 | 16 | 34 |
| OV | Evidence dropout | 0.2 | 0.3 | 0.7 | 50 | 15 | 35 |
| OV | Evidence dropout | 0.4 | 0.3 | 0.7 | 50 | 15 | 35 |
| OV | Evidence dropout | 0.6 | 0.3 | 0.7 | 50 | 15 | 35 |
| OV | Evidence dropout | 0.8 | 0.3 | 0.7 | 50 | 15 | 35 |
| OV | Evidence dropout | 0.9 | 0.24 | 0.76 | 50 | 12 | 38 |
| SKCM | Proposed | 0.2 | 0.66 | 0.34 | 50 | 33 | 17 |
| SKCM | Proposed | 0.4 | 0.66 | 0.34 | 50 | 33 | 17 |
| SKCM | Proposed | 0.6 | 0.66 | 0.34 | 50 | 33 | 17 |
| SKCM | Proposed | 0.8 | 0.66 | 0.34 | 50 | 33 | 17 |
| SKCM | Proposed | 0.9 | 0.62 | 0.38 | 50 | 31 | 19 |
| SKCM | context_swap | 0.2 | 0.42 | 0.58 | 50 | 21 | 29 |

| | | | | | | | |
|---|---|---|---|---|---|---|---|
| SKCM | context_swap | 0.4 | 0.42 | 0.58 | 50 | 21 | 29 |
| SKCM | context_swap | 0.6 | 0.42 | 0.58 | 50 | 21 | 29 |
| SKCM | context_swap | 0.8 | 0.42 | 0.58 | 50 | 21 | 29 |
| SKCM | context_swap | 0.9 | 0.36 | 0.64 | 50 | 18 | 32 |
| SKCM | Evidence dropout | 0.2 | 0.24 | 0.76 | 50 | 12 | 38 |
| SKCM | Evidence dropout | 0.4 | 0.24 | 0.76 | 50 | 12 | 38 |
| SKCM | Evidence dropout | 0.6 | 0.24 | 0.76 | 50 | 12 | 38 |
| SKCM | Evidence dropout | 0.8 | 0.24 | 0.76 | 50 | 12 | 38 |
| SKCM | Evidence dropout | 0.9 | 0.2 | 0.8 | 50 | 10 | 40 |
| UCEC | Proposed | 0.2 | 0.78 | 0.22 | 50 | 39 | 11 |
| UCEC | Proposed | 0.4 | 0.78 | 0.22 | 50 | 39 | 11 |
| UCEC | Proposed | 0.6 | 0.78 | 0.22 | 50 | 39 | 11 |
| UCEC | Proposed | 0.8 | 0.78 | 0.22 | 50 | 39 | 11 |
| UCEC | Proposed | 0.9 | 0.7 | 0.3 | 50 | 35 | 15 |
| UCEC | context_swap | 0.2 | 0.28 | 0.72 | 50 | 14 | 36 |
| UCEC | context_swap | 0.4 | 0.28 | 0.72 | 50 | 14 | 36 |
| UCEC | context_swap | 0.6 | 0.28 | 0.72 | 50 | 14 | 36 |
| UCEC | context_swap | 0.8 | 0.28 | 0.72 | 50 | 14 | 36 |
| UCEC | context_swap | 0.9 | 0.26 | 0.74 | 50 | 13 | 37 |
| UCEC | Evidence dropout | 0.2 | 0.2 | 0.8 | 50 | 10 | 40 |
| UCEC | Evidence dropout | 0.4 | 0.2 | 0.8 | 50 | 10 | 40 |
| UCEC | Evidence dropout | 0.6 | 0.2 | 0.8 | 50 | 10 | 40 |
| UCEC | Evidence dropout | 0.8 | 0.2 | 0.8 | 50 | 10 | 40 |
| UCEC | Evidence dropout | 0.9 | 0.2 | 0.8 | 50 | 10 | 40 |

Proposed (matched Sample Card context), context_swap (shuffled Sample Card context; e.g., BRCA→LUAD), and Evidence dropout (evidence-dropout stress, p = 0.05, min_keep = 1).

**S2.5. Supplementary Table 5 | Human labeling protocol and labeled claims**

Human-labeling protocol for decision-grade acceptability (external endpoint). Part B provides the labeled claim set (claim-level rows) with minimal, ID-centered fields for reproducible evaluation and optional calibration.

**Part A. Human labeling protocol (external endpoint: decision-grade acceptability)**

Human labels reflect decision-grade acceptability under the stated context, not biological mechanistic truth.

**Label set.** Raters assigned one of three labels: **ACCEPT**, **SHOULD_ABSTAIN**, or **REJECT**. **ACCEPT** indicates the claim is acceptable to state as a context-bounded directional enrichment/signature observation (not a mechanistic or causal statement). **SHOULD_ABSTAIN** is reserved for cases where the claim is not safe to communicate as written even under signature framing (e.g., it would require substantial qualifiers to avoid mechanistic implication, or it is not interpretable from the provided context/evidence without guesswork). **REJECT** indicates broken/ambiguous evidence linkage or incompatibility with the stated context.

| question_id | question (yes/no) | if "no" |
|---|---|---|
| Q1 (pre-screen) | Are evidence references complete and unambiguous (all referenced terms/genes/modules can be in the provided EvidenceTable; no broken or ambiguous linkage)? | REJECT |
| Q2 | Is the claim acceptable to state under the stated context (condition/comparison) without overreach (no causal/mechanistic or unwarranted specificity statements)? | REJECT |
| Q3 | Should we abstain because the claim is not safe to communicate as written (i.e., it would require substantial qualifiers to avoid mechanistic implication, or it is not interpretable from the provided context/evidence without guesswork)? | ACCEPT (if "yes" _ SHOULD_ABSTAIN) |

**Part B. Labeled claims (minimum fields)**

Each row represents one claim labeled by one rater. The table contains the minimal fields required to (i) present a context-bounded, evidence-linked item for human judgment and (ii) deterministically link the item back to the audited claim and EvidenceTable-derived artifacts. A distributable labeling template is generated deterministically by paper/scripts/fig2_make_labels_template.py. Provenance-only fields (e.g., _src) may be retained for traceability but are not shown during rating. Required columns: claim_id, human_label, rater_id, label_version, condition, entity, direction, term_name, gene_symbols_str, module_id_effective Optional column: _src

| column | requirement | content / constraints |
|---|---|---|
| claim_id | Core | Stable claim identifier (string). |
| human_label | Core | Human label in {ACCEPT, SHOULD_ABSTAIN, REJECT}. May be empty in the distributed template; only non-empty labels are used for agreement and evaluation. |
| rater_id | Core | Anonymized rater identifier (string). |
| label_version | Core | Labeling protocol version (string; e.g., v1). |
| condition | Core | Condition/cohort label (string; e.g., HNSC, BEATAML). |
| entity | Core | Evidence term identifier referenced by the claim (string; semantically term_id; e.g., HALLMARK_…, GO:…, R-HSA-…). |
| direction | Core | Direction in {up, down, na} (string). |
| term_name | Core | Human-readable term name (string; ≤1 line). |
| gene_symbols_str | Core | Representative evidence genes shown to raters (string; delimiter fixed by export, e.g., ;; ≤1 line). |
| module_id_effective | Core | Effective evidence module identifier used by the auditor (string). |
| _src | (optional) | Provenance pointer to the originating audit_log.tsv (string; for traceability only; not shown during rating). |

**S2.6. Supplementary Table 6 | Reduced audit-log view (LLM-assisted proposal; related to Extended Data Fig. 3).**
This table provides a compact subset of the audit log for the LLM-assisted setting in HNSC (Proposed, $\tau = 0.8$; hard-gated audit; $k = 50$ claims). For each schema-bounded claim, we report the stable claim identifier; the referenced term and direction; the resolved evidence module and supporting-gene set hash; the context-review outcome and the final audit decision (PASS/ABSTAIN/FAIL) together with any associated abstain/fail reason codes and notes. The full, machine-readable audit log (all fields) is provided as Source Data.

| claim_id | term_id | direction | module_id_effective | gene_set_hash_effective | context_status | context_reason | status | abstain_reason | fail_reason | audit_notes |
|---|---|---|---|---|---|---|---|---|---|---|
| c_de0c536ad356 | HALLMARK_APOPTOSIS | up | Mdf6350d89cee | 1b912bf139ec | PASS | LLM_CR1 | PASS | | | |
| c_402f11698cfb | HALLMARK_P53_PATHWAY | up | Maf269dcd4575 | 73cbba12ef1c | PASS | LLM_CR2 | PASS | | | |
| c_53120a9f833e | HALLMARK_TNFA_SIGNALING_VIA_NFKB | up | Mdf6350d89cee | 8f055c2f4077 | PASS | LLM_CR3 | PASS | | | |
| c_07978786a0ce | HALLMARK_ANGIOGENESIS | up | Me6bf443283c2 | e7371f4eac9a | PASS | LLM_CR4 | PASS | | | |
| c_7991fb0c5841 | HALLMARK_WNT_BETA_CATENIN_SIGNALING | up | M591a524fec18 | fdfa5bae0a42 | FAIL | LLM_CR5 | FAIL | | context_fail | AN1 |
| c_6ec10a980027 | HALLMARK_ADIPOGENESIS | up | M03a383e6e086 | ce5fe3ea4387 | FAIL | LLM_CR6 | FAIL | | context_fail | AN2 |
| c_4e1651d119b7 | HALLMARK_APICAL_JUNCTION | up | Mf906280fa5e1 | 37b4810ad81e | PASS | LLM_CR7 | PASS | | | |
| c_287d32df5891 | HALLMARK_HEDGEHOG_SIGNALING | up | M0332a8692e9e | 613ae2fabfbc | PASS | LLM_CR8 | PASS | | | |
| c_3d57869efdf6 | HALLMARK_IL6_JAK_STAT3_SIGNALING | up | Ma4188ca7ae89 | 23229151cd5d | PASS | LLM_CR9 | PASS | | | |
| c_0140522d1577 | HALLMARK_CHOLESTEROL_HOMEOSTASIS | up | M5ff3f8386599 | 01a3aafc0275 | FAIL | LLM_CR10 | FAIL | | context_fail | AN3 |
| c_1f5784486133 | HALLMARK_MITOTIC_SPINDLE | up | M719ba53cf8d5 | 4b5f4290c226 | PASS | LLM_CR11 | PASS | | | |
| c_ad2d4a6f0856 | HALLMARK_TGF_BETA_SIGNALING | up | M6f6e9e05805a | 79fc0935c212 | PASS | LLM_CR12 | PASS | | | |
| c_3128e0bd890b | HALLMARK_UNFOLDED_PROTEIN_RESPONSE | up | Mcf69f9e4157c | 248c1a1dbb2b | PASS | LLM_CR13 | ABSTAIN | unstable | | AN4 |
| c_527e226afd77 | HALLMARK_GLYCOLYSIS | down | Mf4596493a738 | 3477ec8e46f3 | PASS | LLM_CR14 | PASS | | | |

| | | | | | | | | | |
|---|---|---|---|---|---|---|---|---|---|
| c_5e57c66438fe | HALLMARK_HEME_METABOLISM | down | Me43c4cd101bd | 8951f4c8785b | FAIL | LLM_CR15 | FAIL | context_fail | AN5 |
| c_970633c2904e | HALLMARK_ESTROGEN_RESPONSE_EARLY | down | M849a4c0e7f2d | 0d37c176f128 | FAIL | LLM_CR16 | FAIL | context_fail | AN6 |
| c_da0bf28d2afd | HALLMARK_COAGULATION | down | Mc4fe664d694a | 75f6cb08faf0 | PASS | LLM_CR17 | PASS | | |
| c_d9bf3dc81b90 | HALLMARK_NOTCH_SIGNALING | down | Mea4001d70482 | 5c2b02c8d35c | FAIL | LLM_CR18 | FAIL | context_fail | AN7 |
| c_f321e754f77e | HALLMARK_REACTIVE_OXYGEN_SPECIES_PATHWAY | down | Mf1237f2d6d20 | ca3891cda6d4 | FAIL | LLM_CR19 | FAIL | context_fail | AN8 |
| c_c8e37e94d74c | HALLMARK_DNA_REPAIR | down | Mdd3caae7261b | b4c104b803bc | PASS | LLM_CR20 | PASS | | |
| c_025cf777a4d7 | HALLMARK_UV_RESPONSE_DN | down | M029534977e54 | b3127b9c263c | PASS | LLM_CR21 | PASS | | |
| c_69fe7fd74096 | HALLMARK_EPITHELIAL_MESENCHYMAL_TRANSITION | down | Mc4fe664d694a | 92e891fc1bf4 | FAIL | LLM_CR22 | FAIL | context_fail | AN9 |
| c_d94a3ac2ad37 | HALLMARK_PROTEIN_SECRETION | down | Md4f5e0a53b7c | d45d65800549 | FAIL | LLM_CR23 | FAIL | context_fail | AN10 |
| c_1e3c8b7f407f | HALLMARK_MYOGENESIS | down | Me334011e2fa6 | e0341df89c4b | FAIL | LLM_CR24 | FAIL | context_fail | AN11 |
| c_be20f809d56c | HALLMARK_HYPOXIA | down | Mf4596493a738 | aeeeed6d60d8 | PASS | LLM_CR25 | PASS | | |
| c_a3c5ce712645 | HALLMARK_XENOBIOTIC_METABOLISM | down | M557bebe138f3 | f3fe33cde6b1 | FAIL | LLM_CR26 | FAIL | context_fail | AN12 |
| c_1046bdacc5c3 | HALLMARK_PANCREAS_BETA_CELLS | down | M3297d664d230 | 728f7e7d6aee | FAIL | LLM_CR27 | FAIL | context_fail | AN13 |
| c_0ca7b9ffbde8 | HALLMARK_OXIDATIVE_PHOSPHORYLATION | up | M03a383e6e086 | 3fdf9a49257d | PASS | LLM_CR28 | PASS | | |
| c_f2aca0d7fc8f | HALLMARK_ESTROGEN_RESPONSE_LATE | up | M8e5c1d28573e | 0ea081758020 | FAIL | LLM_CR29 | FAIL | context_fail | AN14 |
| c_383845c339a0 | HALLMARK_COMPLEMENT | down | Mc4fe664d694a | f62dd3909e9d | PASS | LLM_CR30 | PASS | | |
| c_0f65c1132227 | HALLMARK_INFLAMMATORY_RESPONSE | down | Mc655fb76a1be | 3888970a3026 | FAIL | LLM_CR31 | FAIL | context_fail | AN15 |
| c_66900b2e124c | HALLMARK_APICAL_SURFACE | down | M7d7e92d830b5 | 4e57c19a9fb1 | FAIL | LLM_CR32 | FAIL | context_fail | AN16 |

| | | | | | | | | | |
|---|---|---|---|---|---|---|---|---|---|
| c_e2d7a94f8047 | HALLMARK_MTORC1_SIGNALING | up | M852823ef4bff | 5bd1e2775dba | PASS | LLM_CR33 | PASS | | |
| c_d96eeb106a52 | HALLMARK_PEROXISOME | up | M316384fde50c | 224b55fc8ce4 | PASS | LLM_CR34 | PASS | | |
| c_b14a39eaf8ab | HALLMARK_INTERFERON_ALPHA_RESPONSE | down | M94039d22fbff | 8cb48550b126 | PASS | LLM_CR35 | PASS | | |
| c_034016a6656c | HALLMARK_FATTY_ACID_METABOLISM | down | M461f65de8e09 | f8bb6661691d | FAIL | LLM_CR36 | FAIL | context_fail | AN17 |
| c_45c5a21822f5 | HALLMARK_PI3K_AKT_MTOR_SIGNALING | down | Ma9ddf8f5a0b5 | 98e9db3f2c37 | PASS | LLM_CR37 | PASS | | |
| c_8cb673d58fc0 | HALLMARK_E2F_TARGETS | down | M0b67a17cb0b6 | 0ad1f44f507a | PASS | LLM_CR38 | PASS | | |
| c_fa7d736c70a7 | HALLMARK_UV_RESPONSE_UP | down | M210904edebae | 9ecb1f267791 | PASS | LLM_CR39 | PASS | | |
| c_ef9dbbf4fb80 | HALLMARK_INTERFERON_GAMMA_RESPONSE | down | M94039d22fbff | 645976544bb2 | FAIL | LLM_CR40 | FAIL | context_fail | AN18 |
| c_adfe198d3fea | HALLMARK_G2M_CHECKPOINT | down | M0b67a17cb0b6 | ff309e380b68 | PASS | LLM_CR41 | PASS | | |
| c_8ebbd59464d3 | HALLMARK_BILE_ACID_METABOLISM | down | M8ff555ff76b4 | 838361fd840f | FAIL | LLM_CR42 | FAIL | context_fail | AN19 |
| c_608ae3cc70b0 | HALLMARK_MYC_TARGETS_V1 | down | M0b67a17cb0b6 | 33c997596369 | FAIL | LLM_CR43 | FAIL | context_fail | AN20 |
| c_05b52ed1a10c | HALLMARK_ALLOGRAFT_REJECTION | down | Me89ae70f36de | fbd15c4dc6a1 | FAIL | LLM_CR44 | FAIL | context_fail | AN21 |
| c_52d565236d64 | HALLMARK_MYC_TARGETS_V2 | down | Me56b74484d20 | 1e36114f6544 | FAIL | LLM_CR45 | FAIL | context_fail | AN22 |
| c_5435f1f06413 | HALLMARK_KRAS_SIGNALING_UP | down | M40cd09ad5a42 | e91565cb32b2 | PASS | LLM_CR46 | PASS | | |
| c_7776423c45c1 | HALLMARK_ANDROGEN_RESPONSE | down | M27669b8fa9e7 | db62abf089f7 | FAIL | LLM_CR47 | FAIL | context_fail | AN23 |
| c_c5847d7415a8 | HALLMARK_KRAS_SIGNALING_DN | down | M740963684598 | 5e7fc4a4dae8 | PASS | LLM_CR48 | PASS | | |
| c_7acfd9e487ae | HALLMARK_SPERMATOGENESIS | down | M63bca3f82c8e | e0abdccfff67 | FAIL | LLM_CR49 | FAIL | context_fail | AN24 |
| c_5e8a60b12983 | HALLMARK_IL2_STAT5_SIGNALING | down | M0b9664e418b9 | bb99dd8b89f9 | PASS | LLM_CR50 | PASS | | |

**Unvalidated LLM-generated rationales (verbatim; not used by audit).**

LLM-generated rationales (LLM_CR*) are provided verbatim for transparency and error analysis and may contain factual errors. In the LLM-assisted proposal mode (used only for **Extended Data Fig. 3**), audit decisions are computed from tool-resolved EvidenceTable links and predefined rule-based gates; free-text rationales are recorded but are not parsed or used by any audit gate.

**llm_rationale_verbatim (context_reason)**

LLM_CR1: The term HALLMARK_APOPTOSIS is relevant to the context of TP53 mutation in a tumor sample (HNSC), as apoptosis is often associated with cancer and TP53 mutation

LLM_CR2: The term HALLMARK_P53_PATHWAY is relevant to the context condition HNSC and perturbation genotype TP53_mut_vs_TP53_wt.

LLM_CR3: The term HALLMARK_TNFA_SIGNALING_VIA_NFKB is associated with genes involved in TNF-alpha signaling via NF-kappa B pathway, which is relevant to the context of H

LLM_CR4: The term HALLMARK_ANGIOGENESIS is associated with angiogenesis, which is a characteristic of tumor tissues.

LLM_CR5: Term 'HALLMARK_WNT_BETA_CATENIN_SIGNALING' does not match context condition 'HNSC'

LLM_CR6: The term HALLMARK_ADIPOGENESIS is not related to the context condition HNSC or tissue tumor.

LLM_CR7: The term HALLMARK_APICAL_JUNCTION is associated with apical junctions in epithelial cells, which are relevant to tumor tissues.

LLM_CR8: The term HALLMARK_HEDGEHOG_SIGNALING is known to be associated with TP53 mutations in the context of HNSC tumors.

LLM_CR9: The term is associated with IL-6/JAK/STAT3 signaling pathway, which is relevant to HNSC (Head and Neck Squamous Cell Carcinoma) and tumor samples.

LLM_CR10: The term HALLMARK_CHOLESTEROL_HOMEOSTASIS is not related to the context condition HNSC or cancer type.

LLM_CR11: The term HALLMARK_MITOTIC_SPINDLE is associated with genes involved in cell cycle regulation, which is consistent with the context of a tumor sample (HNSC) and

LLM_CR12: The term HALLMARK_TGF_BETA_SIGNALING is known to be associated with TGF-beta signaling pathway, which is relevant in the context of TP53 mutation vs wildtype co

LLM_CR13: The term HALLMARK_UNFOLDED_PROTEIN_RESPONSE is associated with cellular stress responses and unfolded protein response pathways are often dysregulated in cancer

LLM_CR14: The term HALLMARK_GLYCOLYSIS is relevant to the context of a tumor (HNSC) with a genotype perturbation involving TP53 mutation.

LLM_CR15: The term 'HALLMARK_HEME_METABOLISM' is not related to the context's condition 'HNSC', which is a specific cancer type.

LLM_CR16: The term 'HALLMARK_ESTROGEN_RESPONSE_EARLY' is not related to the context's condition 'HNSC', which is a type of Head and Neck Squamous Cell Carcinoma, and does

LLM_CR17: The term HALLMARK_COAGULATION is associated with various cancer types and has a high overlap with the given context's signature (context_signature: e6cee1e46cac

LLM_CR18: The term HALLMARK_NOTCH_SIGNALING is not related to the context's condition (HNSC) or comparison (TP53_mut_vs_TP53_wt)

LLM_CR19: The term 'HALLMARK_REACTIVE_OXYGEN_SPECIES_PATHWAY' is not related to the context's condition 'HNSC', which is Head and Neck Squamous Cell Carcinoma, a cancer t

LLM_CR20: The term HALLMARK_DNA_REPAIR is relevant to DNA repair mechanisms, which are often impaired in tumor tissues (like HNSC). The presence of TP53 mutations may als

LLM_CR21: The term HALLMARK_UV_RESPONSE_DN is associated with UV damage response and is relevant to skin cancer (HNSC), which aligns with the context.
LLM_CR22: The term 'HALLMARK_EPITHELIAL_MESENCHYMAL_TRANSITION' does not match the context's condition 'HNSC', which is a specific cancer type, while the term is more gen
LLM_CR23: The term 'HALLMARK_PROTEIN_SECRETION' is not clearly context-consistent with the sample context.
LLM_CR24: The term HALLMARK_MYOGENESIS is not related to the context condition HNSC or tissue tumor.
LLM_CR25: The term HALLMARK_HYPOXIA is associated with hypoxia in various cancer types, including HNSC (Head and Neck Squamous Cell Carcinoma). The context signature e6ce
LLM_CR26: The term 'HALLMARK_XENOBIOTIC_METABOLISM' does not match the context's condition 'HNSC', which is a specific cancer type, whereas the term is related to xenobio
LLM_CR27: Term 'HALLMARK_PANCREAS_BETA_CELLS' is not context-consistent with sample context: condition='HNSC', tissue='tumor'
LLM_CR28: The term HALLMARK_OXIDATIVE_PHOSPHORYLATION is relevant to the context of HNSC tumor samples with TP53 mutation compared to wild-type, as oxidative phosphorylat
LLM_CR29: The term 'HALLMARK_ESTROGEN_RESPONSE_LATE' is not related to the context's condition 'HNSC', which is a type of Head and Neck Squamous Cell Carcinoma, and does
LLM_CR30: The term HALLMARK_COMPLEMENT is associated with various cancer types and has a high overlap with the given context's signature (context_signature: e6cee1e46cac)
LLM_CR31: The term 'HALLMARK_INFLAMMATORY_RESPONSE' is not related to the context's condition 'HNSC', which is a specific cancer type.
LLM_CR32: Term 'HALLMARK_APICAL_SURFACE' is not related to the context's condition 'HNSC'
LLM_CR33: The term HALLMARK_MTORC1_SIGNALING is associated with MTOR signaling pathway, which is relevant to cancer and particularly to TP53 mutations.
LLM_CR34: The term HALLMARK_PEROXISOME is associated with genes involved in peroxisome function, which are relevant to cancer biology and specifically to the context of T
LLM_CR35: The term HALLMARK_INTERFERON_ALPHA_RESPONSE is associated with interferon-alpha signaling pathway, which is relevant to tumor context and TP53 mutation comparis
LLM_CR36: The term HALLMARK_FATTY_ACID_METABOLISM is not related to the context condition HNSC or tissue tumor.
LLM_CR37: The term HALLMARK_PI3K_AKT_MTOR_SIGNALING is associated with PI3K/Akt/mTOR signaling pathway, which is relevant to cancer and particularly to TP53 mutations.
LLM_CR38: The term HALLMARK_E2F_TARGETS is associated with E2F target genes, which are often upregulated in TP53 mutant tumors compared to wild-type tumors.
LLM_CR39: The term HALLMARK_UV_RESPONSE_UP is relevant to skin cancer (HNSC) and involves UV response, which aligns with the context.
LLM_CR40: The term 'HALLMARK_INTERFERON_GAMMA_RESPONSE' is not relevant to the context of HNSC (Head and Neck Squamous Cell Carcinoma) tumor samples, especially consideri
LLM_CR41: The term HALLMARK_G2M_CHECKPOINT is associated with cell cycle regulation and checkpoint control, which is relevant to tumor samples.
LLM_CR42: No clear connection between HALLMARK_BILE_ACID_METABOLISM and the sample context (HNSC tumor)
LLM_CR43: The term 'HALLMARK_MYC_TARGETS_V1' is not expected to be active in a TP53 mutation context.

LLM_CR44: Term 'HALLMARK_ALLOGRAFT_REJECTION' is not context-consistent with sample condition 'HNSC', which is a specific cancer type, whereas the term is related to allo
LLM_CR45: The term 'HALLMARK_MYC_TARGETS_V2' is not clearly context-consistent with the sample context.
LLM_CR46: The term HALLMARK_KRAS_SIGNALING_UP is associated with KRAS signaling pathway, which is relevant to TP53 mutations in HNSC cancer type.
LLM_CR47: The term 'HALLMARK_ANDROGEN_RESPONSE' is not related to the context's condition 'HNSC', which is Head and Neck Squamous Cell Carcinoma, a cancer type that does
LLM_CR48: The term HALLMARK_KRAS_SIGNALING_DN is associated with KRAS mutations, which are relevant to the TP53_mut_vs_TP53_wt comparison in the context of HNSC tumors.
LLM_CR49: Term HALLMARK_SPERMATOGENESIS is not context-consistent with sample condition HNSC and tissue tumor.
LLM_CR50: The term HALLMARK_IL2_STAT5_SIGNALING is associated with IL-2 signaling pathway, which is relevant to tumor samples and TP53 mutations

### audit_notes

AN1: context_fail(hard): context_evaluated=True method=llm status=FAIL reason=Term 'HALLMARK_WNT_BETA_CATENIN_SIGNALING' does not match context condition 'HNSC'
AN2: context_fail(hard): context_evaluated=True method=llm status=FAIL reason=The term HALLMARK_ADIPOGENESIS is not related to the context condition HNSC or tissue tumor.
AN3: context_fail(hard): context_evaluated=True method=llm status=FAIL reason=The term HALLMARK_CHOLESTEROL_HOMEOSTASIS is not related to the context condition HNSC or cancer type.
AN4: survival[term]=0.797 < tau=0.80
AN5: context_fail(hard): context_evaluated=True method=llm status=FAIL reason=The term 'HALLMARK_HEME_METABOLISM' is not related to the context's condition 'HNSC', which is a specific cancer type.
AN6: context_fail(hard): context_evaluated=True method=llm status=FAIL reason=The term 'HALLMARK_ESTROGEN_RESPONSE_EARLY' is not related to the context's condition 'HNSC', which is a type of Head and Neck Squamous Cell Carcinoma, and does
AN7: context_fail(hard): context_evaluated=True method=llm status=FAIL reason=The term HALLMARK_NOTCH_SIGNALING is not related to the context's condition (HNSC) or comparison (TP53_mut_vs_TP53_wt)
AN8: context_fail(hard): context_evaluated=True method=llm status=FAIL reason=The term 'HALLMARK_REACTIVE_OXYGEN_SPECIES_PATHWAY' is not related to the context's condition 'HNSC', which is Head and Neck Squamous Cell Carcinoma, a cancer t
AN9: context_fail(hard): context_evaluated=True method=llm status=FAIL reason=The term 'HALLMARK_EPITHELIAL_MESENCHYMAL_TRANSITION' does not match the context's condition 'HNSC', which is a specific cancer type, while the term is more gen
AN10: context_fail(hard): context_evaluated=True method=llm status=FAIL reason=The term 'HALLMARK_PROTEIN_SECRETION' is not clearly context-consistent with the sample context.
AN11: context_fail(hard): context_evaluated=True method=llm status=FAIL reason=The term HALLMARK_MYOGENESIS is not related to the context condition HNSC or tissue tumor.
AN12: context_fail(hard): context_evaluated=True method=llm status=FAIL reason=The term 'HALLMARK_XENOBIOTIC_METABOLISM' does not match the context's condition 'HNSC', which is a specific cancer type, whereas the term is related to xenobio

AN13: context_fail(hard): context_evaluated=True method=llm status=FAIL reason=Term 'HALLMARK_PANCREAS_BETA_CELLS' is not context-consistent with sample context: condition='HNSC', tissue='tumor'
AN14: context_fail(hard): context_evaluated=True method=llm status=FAIL reason=The term 'HALLMARK_ESTROGEN_RESPONSE_LATE' is not related to the context's condition 'HNSC', which is a type of Head and Neck Squamous Cell Carcinoma, and does
AN15: context_fail(hard): context_evaluated=True method=llm status=FAIL reason=The term 'HALLMARK_INFLAMMATORY_RESPONSE' is not related to the context's condition 'HNSC', which is a specific cancer type.
AN16: context_fail(hard): context_evaluated=True method=llm status=FAIL reason=Term 'HALLMARK_APICAL_SURFACE' is not related to the context's condition 'HNSC'
AN17: context_fail(hard): context_evaluated=True method=llm status=FAIL reason=The term HALLMARK_FATTY_ACID_METABOLISM is not related to the context condition HNSC or tissue tumor.
AN18: context_fail(hard): context_evaluated=True method=llm status=FAIL reason=The term 'HALLMARK_INTERFERON_GAMMA_RESPONSE' is not relevant to the context of HNSC (Head and Neck Squamous Cell Carcinoma) tumor samples, especially consideri
AN19: context_fail(hard): context_evaluated=True method=llm status=FAIL reason=No clear connection between HALLMARK_BILE_ACID_METABOLISM and the sample context (HNSC tumor)
AN20: context_fail(hard): context_evaluated=True method=llm status=FAIL reason=The term 'HALLMARK_MYC_TARGETS_V1' is not expected to be active in a TP53 mutation context.
AN21: context_fail(hard): context_evaluated=True method=llm status=FAIL reason=Term 'HALLMARK_ALLOGRAFT_REJECTION' is not context-consistent with sample condition 'HNSC', which is a specific cancer type, whereas the term is related to allo
AN22: context_fail(hard): context_evaluated=True method=llm status=FAIL reason=The term 'HALLMARK_MYC_TARGETS_V2' is not clearly context-consistent with the sample context.
AN23: context_fail(hard): context_evaluated=True method=llm status=FAIL reason=The term 'HALLMARK_ANDROGEN_RESPONSE' is not related to the context's condition 'HNSC', which is Head and Neck Squamous Cell Carcinoma, a cancer type that does
AN24: context_fail(hard): context_evaluated=True method=llm status=FAIL reason=Term HALLMARK_SPERMATOGENESIS is not context-consistent with sample condition HNSC and tissue tumor.
